# Taste evolution in an herbivorous drosophilid

**DOI:** 10.1101/2024.02.27.582299

**Authors:** Julianne N. Peláez, Susan Bernstein, Judith Okoro, Esteban Rodas, Irene Liang, Anna Leipertz, Frédéric Marion-Poll, Noah K. Whiteman

**Affiliations:** Department of Integrative Biology, University of California-Berkeley, Berkeley, CA 94720, USA; Department of Molecular & Cellular Biology, University of California-Berkeley, Berkeley, CA 94720, USA; Evolution, Genomes, Behaviour and Ecology, IDEEV, CNRS, Université Paris-Saclay, IRD, Gif-sur-Yvette, France; Université Paris-Saclay, AgroParisTech, 91120 Palaiseau, France

## Abstract

Plant secondary metabolites pose a challenge for generalist herbivorous insects because they are not only potentially toxic, they also may trigger aversion. On the contrary, some highly specialized herbivorous insects evolved to use these same compounds as ‘token stimuli’ for unambiguous determination of their host plants. Two questions that emerge from these observations are how recently derived herbivores evolve to overcome this aversion to plant secondary metabolites and the extent to which they evolve increased attraction to these same compounds. In this study, we addressed these questions by focusing on the evolution of bitter taste preferences in the herbivorous drosophilid *Scaptomyza flava*, which is phylogenetically nested deep in the paraphyletic *Drosophila*. We measured behavioral and neural responses of *S. flava* and a set of non-herbivorous species representing a phylogenetic gradient (*S. pallida, S. hsui*, and *D. melanogaster*) towards host- and non-host derived bitter plant compounds. We observed that *S. flava* evolved a shift in bitter detection, rather than a narrow shift towards glucosinolates, the precursors of mustard-specific defense compounds. In a dye-based consumption assay, *S. flava* exhibited shifts in aversion toward the non-mustard bitter, plant-produced alkaloids caffeine and lobeline, and reduced aversion towards glucosinolates, whereas the non-herbivorous species each showed strong aversion to all bitter compounds tested. We then examined whether these changes in bitter preferences of *S. flava* could be explained by changes in sensitivity in the peripheral nervous system and compared electrophysiological responses from the labellar sensilla of *S. flava*, *S. pallida*, and *D. melanogaster*. Using scanning electron microscopy, we also created a map of labellar sensilla in *S. flava* and *S. pallida*. We assigned each sensillum to a functional sensilla class based on their morphology and initial response profiles to bitter and sweet compounds. Despite a high degree of conservation in the morphology and spatial placement of sensilla between *S. flava* and *S. pallida*, electrophysiological studies revealed that *S. flava* had reduced sensitivity to glucosinolates to varying degrees. We found this reduction only in I type sensilla. Finally, we speculate on the potential role that evolutionary genetic changes in gustatory receptors between *S. pallida* and *S. flava* may play in driving these patterns. Specifically, we hypothesize that the evolution of bitter receptors expressed in I type sensilla may have driven the reduced sensitivity observed in *S. flava*, and ultimately, its reduced bitter aversion. The *S. flava* system showcases the importance of reduced aversion to bitter defense compounds in relatively young herbivorous lineages, and how this may be achieved at the molecular and physiological level.

## INTRODUCTION

The evolution of dietary shifts in animals relies on the ability to sense novel chemicals and textures in food and decide whether to consume it. This can be observed in species as diverse as octopi that search the seafloor, tasting with their tentacles (Wells, 1963), to newborn human and non-human primates, that wince at the taste of bitter quinine (Steiner et al., 2001). In terrestrial animals, these sensations often involve not only long-range attraction mediated by vision and olfaction, but also short-range tactile gustation (taste).

Taste has been less well-studied in the context of dietary shifts but operates as a final checkpoint for the acceptance or rejection of the substrate as a food source for an animal or its offspring (Scott, 2018). The evolution of dietary shifts is associated with the dynamic evolution of large families of chemosensory proteins expressed in chemosensory organs (Edger et al., 2015; McBride, 2007). However, what remains unclear are the nature of steps through which feeding preferences evolve, the mechanisms and sensory cues involved, and how genetic changes in chemosensory genes translate to feeding choices across species with different diets.

In no other group has chemosensory evolution been studied more than among herbivorous insects, which account for half of all insects and a quarter of all known eukaryotic species (Bernays, 1998; Wiens et al., 2015). A critical interface between herbivorous insects and their host plants are plant secondary metabolites, many of which have evolved and diversified in a chemical arms race and repel and poison would-be herbivores (Ehrlich & Raven, 1964).

Plant secondary metabolites pose a challenge for insects to evolve herbivory because they are not only potentially toxic to non-specialist insects, they also may trigger an innate aversion response. Studies on gustatory evolution in generalist herbivores - insects feeding on a broad range of plant families - have illuminated a strong association between host breadth and gustatory receptors (GR) diversity: specifically, dramatic expansions of GR repertoires have been implicated in driving the detection of a large diversity of host plant species, given the differential ligand specificity of different GRs (Briscoe et al., 2013; Smadja et al., 2009; Suzuki et al., 2018; Wanner & Robertson, 2008; Xu et al., 2016).

However, most herbivores, over 80%, are oligophagous and feed on one or a restricted set of plant families (Schoonhoven & Jermy, 1998). In contrast to generalist herbivores, much of the literature on taste preferences of specialists has focused on attraction by herbivores to compounds derived from their host plant that act as stimulants or host identifiers. Because of their specialization on one or a few plant families, these ‘token stimuli’ provide unambiguous identification of their hosts (Lipke & Fraenkel, 1956; Verschaffelt, 1910). For example, Brassicales-derived glucosinolates strongly stimulate feeding and oviposition in *Pieris* butterfly species (Chew & Renwick, 1995). In several specialist herbivores, individual GRs that detect these plant compounds have even been identified: Chinese citrus fly *Bactrocera minax* uses *BminGr59f* to detect compounds from unripe citrus fruits (Zhang et al., 2022); *Pieris rapae* uses *PrapGr28* to detect the glucosinolate sinigrin from their host plants (Yang et al., 2021); and *Papilio xuthus,* a specialist on Rutaceae, uses *PxutGr1* to detect synephrine, an oviposition stimulant (Ozaki et al., 2011). However, most herbivores studied first evolved to become herbivorous over 100 million years ago (Gloss et al., 2019), and so we have little insight into gustatory evolution during the early stages of herbivory and host plant specialization.

Examples from *Drosophila* include more recently evolved specialists, like *D. sechellia* and *D. yakuba mayottensis* (monophagous noni specialists) and *D. suzukii* (polyphagous ripe fruit specialist). In response to feeding on bitter hosts, both *D. sechellia* and *D. suzukii* have experienced an overall evolutionary loss of aversion to bitter compounds, likely due to dramatic gene losses and/or down-regulation of GRs tuned to bitter compounds (H. K. Dweck et al., 2021; H. K. M. Dweck & Carlson, 2020; McBride et al., 2007). In addition, *D. sechellia* and *D. yakuba mayottensis* have evolved increased sensitivity and attraction to specific host plant compounds due to evolutionary changes in various chemoreceptors (Dekker et al., 2006; Ferreira et al., 2020; Yassin et al., 2016). However, with few examples of recently evolved herbivore lineages, particularly foliar herbivores, it remains unclear whether host plant specific compounds evolve as feeding stimulants early on in the transition to herbivory, and whether specialists evolve greater or lower sensitivity to bitter compounds (Bernays et al., 2000; Bernays & Chapman, 1994; Schoonhoven & Van Loon, 2002).

To address this gap in our knowledge, we investigated gustatory evolution in the herbivore *Scaptomyza flava* (**Fig. 1a**), which is a member of an herbivorous subgenus within the genus Scaptomyza. The Scaptomyza genus forms a clade sister to the Hawaiian *Drosophila* and phylogenetically nested within the paraphyletic *Drosophila* genus (Lapoint et al., 2013). Most herbivores from this clade are specialists on mustard plants in the Brassicaceae family and relatives in the Brassicales. *S. flava* offers a useful context to investigate bitter taste evolution because this species can be readily caught from the wild and reared in the lab on the model plant *Arabidopsis thaliana* (*Whiteman et al., 2011*). Close non-herbivore relatives, like *S. pallida* and *S. hsui*, each from distinct but closely related subgenera *Parascaptomyza* and *Hemiscaptomyza*, respectively can also be easily reared in the lab. Genome assemblies for these Scaptomyza species and many other drosophilid species (Kim et al., 2021), which, alongside decades of accumulated knowledge on the molecular and neural basis of *D. melanogasters* gustatory system, renders *S. flava* as a prime system to dissect bitter taste evolution.

**Figure 1.**
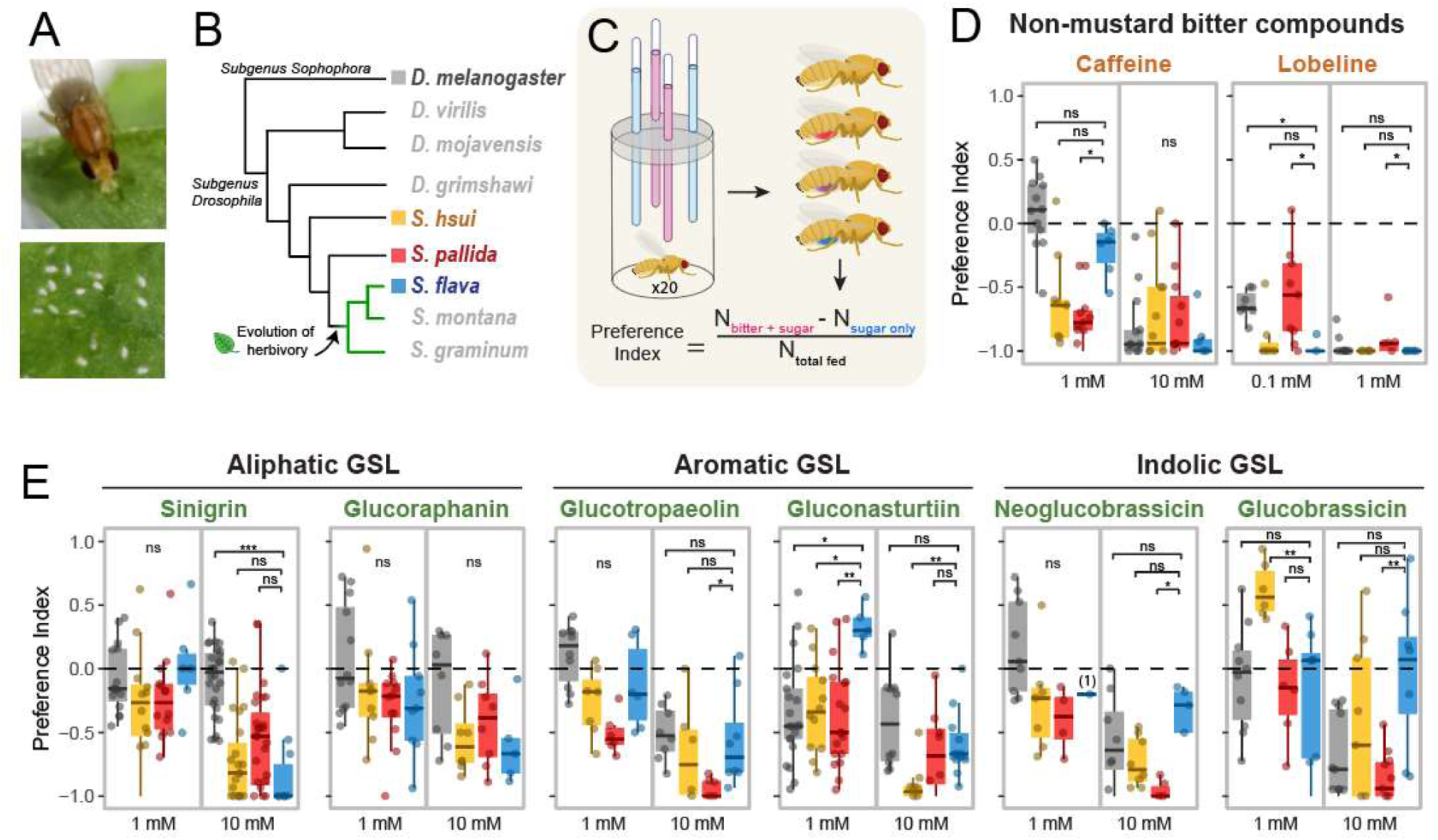
*Scaptomyza flava*, am herbivorous specialist on mustard plants, shows reduced bitter aversion to plant-derived compounds. **(a)** Above, a female *S. flava* feeds on leaf exudates that seeped into wounds created by her serrated ovipositor. Below, a female has laid eggs into similar leaf wounds. **(b)** Phylogenetic placement of the herbivore *S. flava* among microbe-feeding drosophilids. **(c)** Schematic of the two-choice feeding assay. Feeding preferences towards **(d)** non-mustard bitter compounds (caffeine and lobeline) and **(e)** mustard-specific glucosinolates, the precursor molecules to toxic isothiocyanates. Glucosinolates are grouped by class based on the amino acids they are derived from: aromatic (phenylalanine, tyrosine), indolic (tryptophan), and aliphatic (methionine). For panels b and c, bar graphs represent the average, and error bars represent the standard error of the mean. PI = 0 indicates no preference towards the bitter tastant, PI < 0 indicates aversion, and PI > 0 indicates attraction. Preference differences among species were tested with Kruskal-Wallis test, followed by Post-hoc Dunn’s test (*p<0.05, **p<0.01, ***p<0.001, ns = not significant). For panels c and d, n ≥ 5 vials of 10­20 flies, unless specified in parentheses.

The primary chemical defenses in the Brassicales are derived from glucosinolates, sulfur-rich anionic thioglucosidases that are hydrolyzed by the plant enzyme myrosinase (a beta glucosidase) upon tissue damage (Halkier & Gershenzon, 2006). The hydrolysis of glucosinolates by myrosinases - the so-called “mustard oil bomb” - generates several different breakdown products (e.g. isothiocyanates, thiocyanates, and nitriles). Isothiocyanates are toxic to insects because of their strong electrophilic reactivity, binding to nucleic acids and cysteine and lysine residues in proteins and cleaving disulfide bonds (Kawakishi et al., 1983). Despite the toxicity of their breakdown products, glucosinolates are used by more than 25 herbivorous mustard specialists (e.g., the cabbage white *Pieris rapae* and cabbage root fly *Delia radicum*) as token stimuli (Fahey et al., 2001; Hopkins et al., 2008). These insects use glucosinolates, which are largely hydrophilic and non-toxic, as cues to stimulate feeding and egg-laying. However, responses to individual glucosinolates vary depending on the particular glucosinolate, glucosinolate class, and insect species (Bidart-Bouzat & Kliebenstein, 2008).

Glucosinolates are categorized into one of three major structural groups based on the amino acids from which their side chain is derived: aliphatic glucosinolates mainly from methionine, indole glucosinolates from tryptophan, and aromatic glucosinolates from phenylalanine or tyrosine. Not all mustard specialists exhibit the same pattern of responses to these different classes, but there are some general trends. Specialists tend to be indifferent towards indolic glucosinolates. The mustard oils they form are unstable, breaking down quickly into other derivative, potentially less toxic products like indole-3 carbinol (Wittstock et al. 2003). In contrast, many specialists have retained sensitivity towards aliphatics, which produce more toxic and stable mustard oils, which can reduce herbivore fitness (Kliebenstein et al. 2004). Because *S. flava* has acquired an attraction towards the breakdown products of glucosinolates including isothiocyanates (Matsunaga et al., 2022), we hypothesized that in addition to lost aversion to glucosinolates, they might have also gained attraction towards these toxin precursors for increased efficiency in finding appropriate hosts.

To gain insight into changes in the gustatory system of relatively recently evolved specialist herbivores, we examined the behavioral and neural responses of *S. flava* towards plant bitter compounds, contrasting its responses against those of non-herbivorous close relatives that represent a phylogenetic gradient in relatedness with respect to *S. flava*: *S. pallida*, *S. hsui*, and *D. melanogaster*. We utilized a consumption-based feeding assay to identify changing patterns of general bitter detection and changes towards mustard-specific compounds. We then used scanning electron microscopy (SEM) and single sensillum electrophysiology to investigate whether there were alterations at the level of labellar sensilla sensitivity that could explain the behavioral differences we observed. Altogether, we identified a mechanism by which glucosinolate sensitivity was reduced in *S. flava*, while retaining some general bitter sensitivity, and speculate on how evolutionary genetic changes among candidate gustatory receptors might have driven these changes.

## MATERIALS and METHODS

### Fly stocks and maintenance

The species we studied has different dietary niches, which required different rearing conditions and larval diet. *D. melanogaster* (wild-type Canton-S) were maintained on standard Bloomington fly media, provided by the UC Berkeley fly food facility. A colony of *S. flava* (originally collected near Dover, NH, USA) was maintained on *Arabidopsis thaliana* (Col-0 accession) and 10% honey-water solution. Colonies of *S. pallida* and *S. hsui* (bred from isofemale lines, collected from Berkeley, CA, USA) were reared on Bloomington media topped with previously frozen, organic chopped spinach. All fly species were maintained at 22°C ± 2°C, 12L/12D photoperiod, and 70% relative humidity.

Only three species - S. *flava, *S. pallida*,* and *D. melanogaster - were* used for electrophysiological recordings for practical reasons. For these experiments, *S. pallida* and *D. melanogaster* were reared on Nutri-Fly German food formulation (Genessee, Cat. No. 66-115), while *S. flava* was reared on *A. thaliana* (Col-0).

### Feeding assay

To test the feeding preferences of each species, we used a two-choice feeding assay. Newly emerged adult flies were provided honey water *ad libitum*. Roughly 10-20 mated female flies (5-8 days old) were transferred to a vial with moistened filter paper and starved at room temperature for 24 hours. Flies were briefly knocked out with CO_2_ and transferred to the experimental vials. These vials were capped with 3D-printed lids through which four capillary tubes were inserted (**Fig. 1b**). The capillaries had alternating solutions of 5mM sucrose alone and 5mM sucrose with a bitter tastant (0.1, 1, or 10mM) (**Fig. 1b**). Red (0.2 mg/ml sulforhodamine B) and blue (0.08 mg/ml erioglaucine) dyes were added to the two solutions, and were randomly assigned to either solution across different experiments to minimize any effect of dye preferences (Mack & Zhang, 2021). To characterize how herbivorous species may differ from their non-herbivorous relatives in their breadth and sensitivity towards bitter tastants, the test compounds included both those that have been found to elicit aversive responses in *D. melanogaster*, as well as compounds found in mustard plants, specifically glucosinolates, the precursor molecules to the toxic isothiocyanates (listed in Table 1). After feeding in the dark for nine hours, flies were anesthetized with CO_2_ and abdomens were scored as red, blue, purple, or no color. In pilot feeding assays, nine hours was determined as an optimal run time to allow for a majority of *S. flava* and *S. hsui* to feed. However, a limitation of this length of time is that we cannot control for pre- and pos-ingestive effects. Five replicates (vials) were performed for each species per tastant at a given concentration, unless specified otherwise.

**Table 1.**
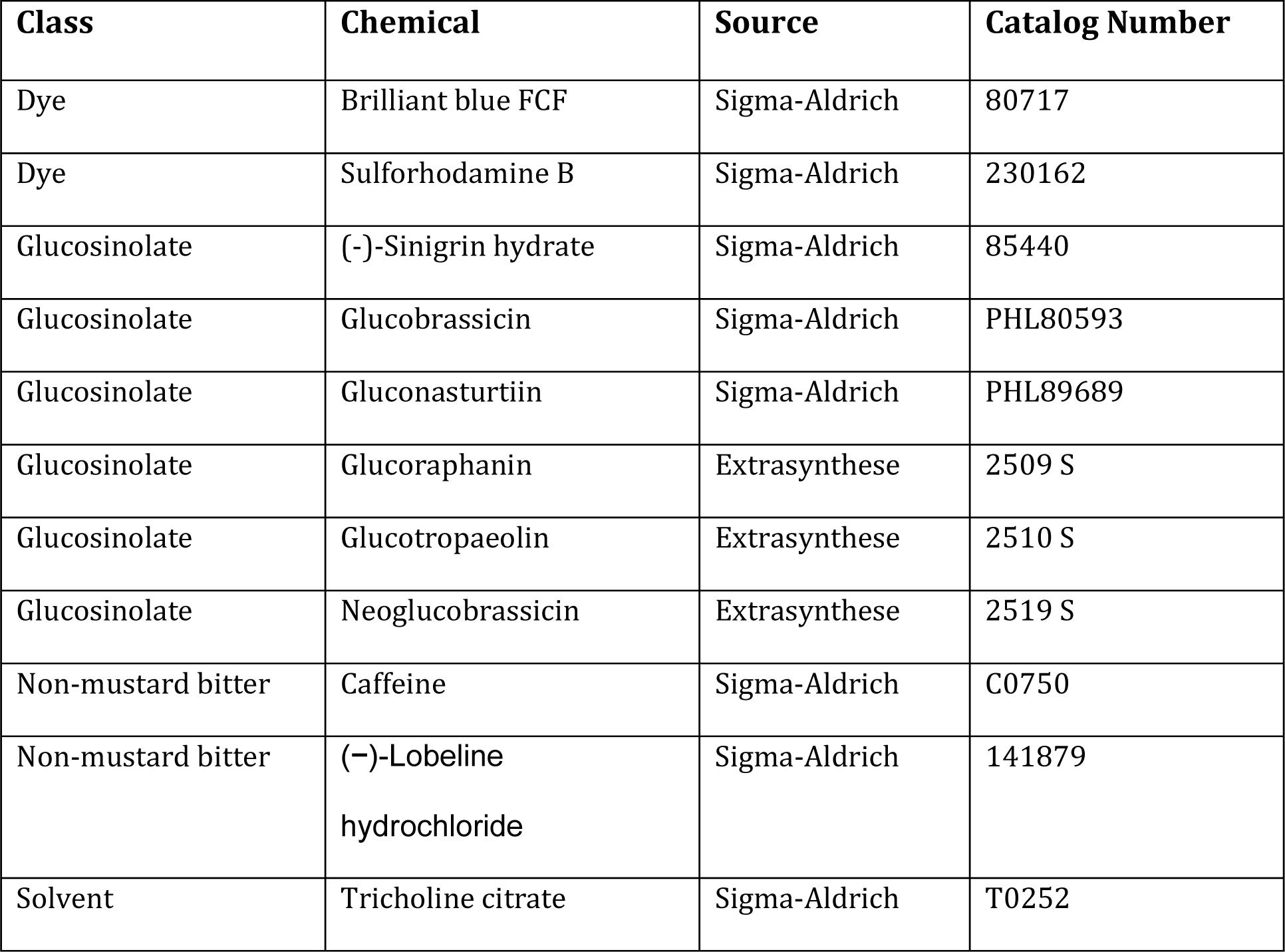
Chemical compounds used in feeding experiments and electrophysiology

| Class | Chemical | Source | Catalog Number |
| --- | --- | --- | --- |
| Dye | Brilliant blue FCF | Sigma-Aldrich | 80717 |
| Dye | Sulforhodamine B | Sigma-Aldrich | 230162 |
| Glucosinolate | (-)-Sinigrin hydrate | Sigma-Aldrich | 85440 |
| Glucosinolate | Glucobrassicin | Sigma-Aldrich | PHL80593 |
| Glucosinolate | Gluconasturtiin | Sigma-Aldrich | PHL89689 |
| Glucosinolate | Glucoraphanin | Extrasynthese | 2509 S |
| Glucosinolate | Glucotropaeolin | Extrasynthese | 2510 S |
| Glucosinolate | Neoglucobrassicin | Extrasynthese | 2519 S |
| Non-mustard bitter | Caffeine | Sigma-Aldrich | C0750 |
| Non-mustard bitter | (-)-Lobeline hydrochloride | Sigma-Aldrich | 141879 |
| Solvent | Tricholine citrate | Sigma-Aldrich | T0252 |

For each tastant, experiments were performed across at least three separate days. A preference index (PI) was calculated for each vial of flies with PI = (number of flies with red abdomens - the number of flies with blue abdomens)/(total number of flies with any color in their abdomens), if the red dye was added to the bitter solution (vice versa if blue dye was added). Preference data were analyzed for differences between species using the Kruskal-Wallis test in R v4.2.1. If significant differences were found (P < 0.05), post-hoc Dunn’s tests were performed to determine which species had significantly different preferences.

### Scanning electron microscopy (SEM)

To identify sensillum types based on morphology among the labellar sensilla in *S. pallida* and *S. flava*, SEM was performed on adult female flies of each species. Fly heads were separated from their bodies to improve the penetration of fixatives into the heads. Heads were fixed in 2% glutaraldehyde in 0.1M sodium cacodylate, stained with osmium tetroxide, and dried through an ethanol series (35-100%). The ethanol was evaporated with a critical point dryer (Tousimis AutoSamdri 815). Specimens were mounted onto stubs, coated with gold-palladium using a Tousimis sputter coater, and imaged with a Hitachi SM-5000 scanning electron microscope (200 - 1500x). All SEM work was carried out at the UC Berkeley Electron Microscope Laboratory.

### Electrophysiology

Newly eclosed flies were transferred to German food formulation for several days, allowed to mate, and then starved on water for 24 hours prior to recordings. Females (5-10 days old) were immobilized and prepared as previously described (French et al., 2015). Using the tip recording method (Hodgson & Roeder, 1956), the recording electrode simultaneously delivers a tastant and transmits an electrical signal to a TasteProbe amplifier (Marion-Poll & van der Pers, 1996). The electric signal was further amplified, filtered (100-3000 Hz; CyberAmp 320) and digitally sampled at 10 kHz with 16 bits precision during 2 s epochs (DT9800; Data Translation). The recordings were acquired and analyzed with the custom software dbWave. Spikes were manually sorted, based on the spike amplitude and shape.

Tricholine citrate (TCC, 30mM) was added to all solutions to silence the water cell (Wieczorek & Wolff, 1989). Freshly made solutions were used for only one week. For each tastant, 5-15 recordings were made per sensilla. Stimulations to the same sensilla were separated at least by ten minutes.

We first performed recordings from all labellar sensilla using 10 mM sucrose, 1 mM lobeline, and 10 mM caffeine to determine whether *S. flava* and *S. pallida* had similar sensillar classes as *D. melanogaster*. In *D. melanogaster*, these three compounds can be used to differentiate L, I-a, I-b, S-a, S-b, and S-c classes of labellar sensilla (Weiss et al., 2011). Because behavioral responses to these compounds were similar between Scaptomyza species and *D. melanogaster*, this suggested that sensillar sensitivity to these compounds might be evolutionarily conserved. We then used a hierarchical cluster analysis applying Ward’s classification method in the PAST program to identify distinct sensillar classes based on their responses to these three compounds (Hammer et al., 2001). Based on clustering, we chose representative sensilla from each class. To further test the differences in sensitivity of these sensilla, we generated a dose response curve for lobeline (0.1, 1, 10 mM) and sucrose and caffeine (1, 10, 100 mM).

Recordings were then performed using glucosinolates (listed in Table 1). Gluconasturtiin, neoglucobrassicin, and sinigrin were tested at 0.1, 1, 10, and 30mM concentrations, and glucoraphanin, glucobrassicin, and glucoraphanin were tested at 1 and 10mM concentrations.

### Gustatory receptor evolution

To identify putative genetic changes that are associated and may even underlie behavioral and physiological differences between herbivorous and non-herbivorous species, we considered gene copy number changes and signatures of positive selection on GR genes. We focused on GRs because of their role in detecting bitter compounds within afferent neurons in the proboscis and tarsi. We have previously reported copy number data and tests for selection on GRs across *Drosophila* and Scaptomyza (Pelaez et al. 2022). This included two non-herbivorous Scaptomyza species (*S. pallida* and S. *hsui)* and three herbivorous *Scaptomyza* species - two of which are leaf-mining specialists on Brassicaceae (*S. flava* and *S. montana*) and the third on Caryophyllaceae (*S. graminum*). Without additional behavioral or physiological data from these additional herbivorous species, we considered any genetic changes shared by all three herbivores, by both mustard feeders, or those only in *S. flava* as genetic candidates to explain the differences among *S. flava*, *S. pallida*, and *S. hsui*.

Previously, we performed selection tests using Phylogenetic Analysis by Maximum Likelihood (PAML), but only reported tests in which the herbivorous clade was specified as the foreground branch (Pelaez et al. 2022). Here, following the same methods, we also ran additional branch-site tests specifying either the two mustard feeders or *S. flava* as the foreground.

## RESULTS

### Mustard specialist shows less aversion to some bitter compounds compared to microbe-feeders

All species, including the herbivore *S. flava*, were averse to the non-mustard specific compounds caffeine and lobeline (**Fig. 1c**). However, at a lower concentration of 1mM caffeine, *S. flava* experienced reduced aversion compared to the non-herbivorous Scaptomyza (*S. flava*, PI=0.4; *S. pallida*, PI=0.14; *S. hsui*, PI=0.19). In contrast, while most other microbe-feeders showed less aversion to lobeline at 1mM compared to at 10 mM, *S. flava* showed almost complete aversion even at 1mM (PI=0.01), showing stronger aversion than *S. pallida* (PI=0.22), although the same level as *S. hsui* (PI=0.05). These results indicate that while *S. flava* has evolved weaker aversion to some bitter compounds (caffeine), they may have evolved stronger aversion towards others (lobeline).

The responses towards glucosinolates varied depending on the species and glucosinolate class (**Fig. 1d**). Most glucosinolates elicited aversive responses in all species. The strongest aversive responses were from aromatic and indolic glucosinolates, with *S. pallida* showing almost complete aversion towards some compounds (e.g. 10 mM glucotropaeolin, PI=0.03; 10 mM neoglucobrassicin, PI=0.02). However, *S. flava* found aromatic glucosinolates (glucotropaeolin and gluconasturtiin) less aversive compared to non-herbivorous Scaptomyza. *S. flava* showed no preference at the lower concentration of 1 mM (glucotropaeolin, PI=0.43; gluconasturtiin, PI=0.66). In response to indolic glucosinolates (neoglucobrassicin and glucobrassicin), *S. flava* showed almost complete indifference even at a higher concentration of 10 mM (neoglucobrassicin, PI=0.4; glucobrassicin, PI=0.48), relative to non-herbivorous Scaptomyza (i.e., *S. pallida*: neoglucobrassicin, PI=0.02; glucobrassicin, PI=0.08).

All species were generally more indifferent towards aliphatic glucosinolates (sinigrin and glucoraphanin) than those towards aromatic and indolic glucosinolates. Furthermore, we did not find statistically significant reductions in aversion towards glucoraphanin in *S. flava* at either of the concentrations tested (1 and 10mM), and only reduced aversion towards sinigrin at 10mM. Interestingly, *D. melanogaster* did not show any aversion or preference towards the aliphatic glucosinolates at all (Pl≈0.5 across all tested concentrations of both aliphatic compounds), but in general, *D. melanogaster* also showed lower aversion towards all the other glucosinolates, relative to the non-herbivorous Scaptomyza. Overall, these behavioral experiments involving glucosinolates suggest that *S. flava* evolved reduced aversion to glucosinolates, but preferences varied across different glucosinolates in herbivores and non-herbivores alike.

### Conservation of sensilla number and location within Scaptomyza

Using scanning electron microscopy, we found that the number and location of sensilla were largely conserved between *D. melanogaster* and Scaptomyza species, and even more similar between *S. flava* and *S. pallida* (**Fig. 2a**). All three species shared the same number and location of long L type sensilla. There were, however, typically two fewer intermediate I type sensilla along the dorsal region of the labellum in both *S. flava* and *S. pallida*, compared to *D. melanogaster*. Similarly, there was a slight difference in S type sensilla between the two genera, with 12 S type sensilla found in the two Scaptomyza species, and typically 10 in *D. melanogaster*. The number of I and S type sensilla varied across individuals of *D. melanogaster* as well (Hiroi et al., 2002). We observed during electrophysiological recordings that both I and S type sensilla numbers were variable across all 3 species, with either one more or fewer present across some individuals. It is unclear the extent to which this lower number of I type sensilla and higher number of S type in Scaptomyza impacts their taste preferences. However, these comparisons clearly show that between Scaptomyza species, the number and location of sensilla types are evolutionarily conserved, which suggests individual sensilla between *S. flava* and *S. pallida* are homologous and their electrophysiology sensitivities can therefore be compared.

**Figure 2.**
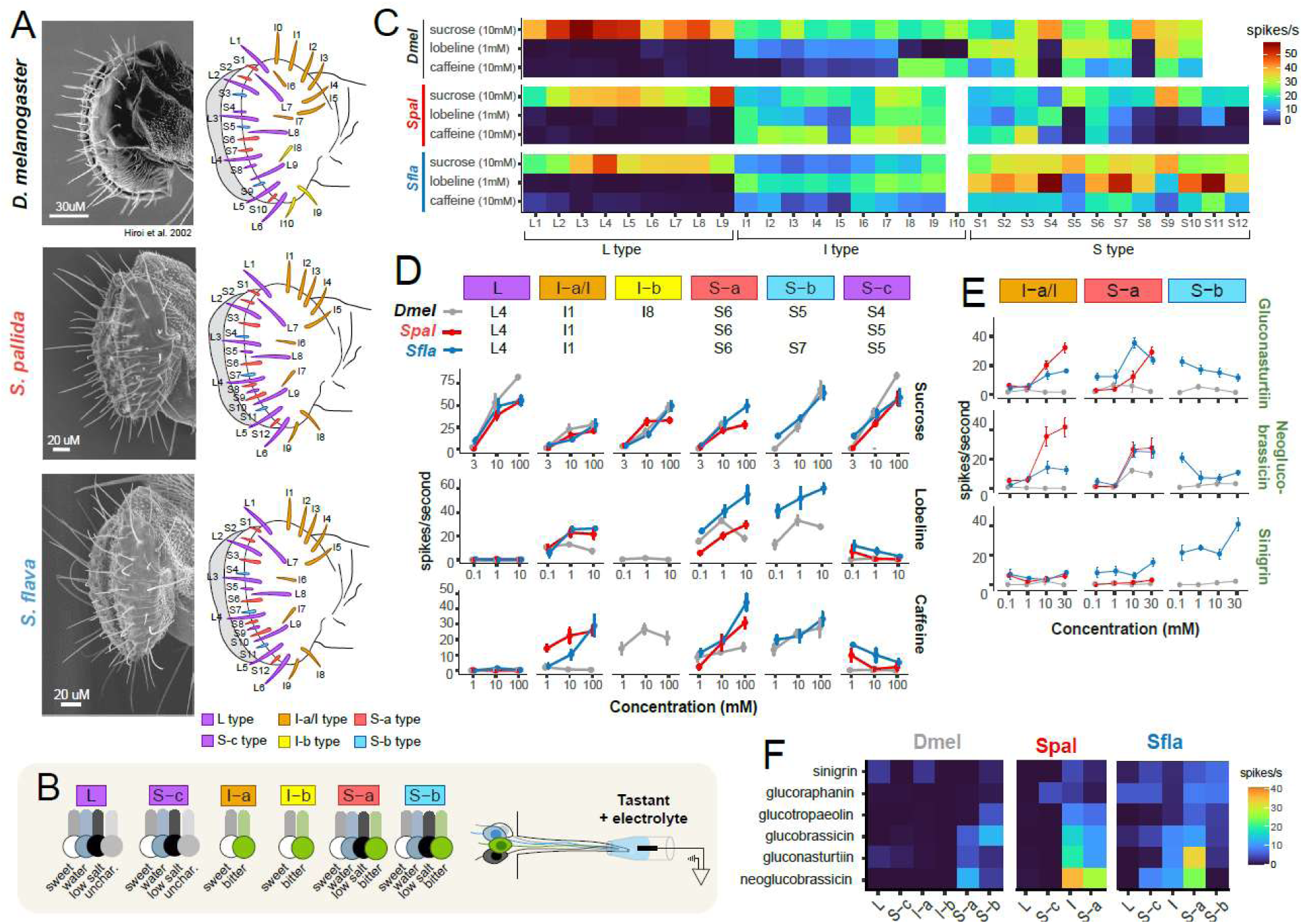
Labellar sensilla types are conserved across *D. melanogaster*, *S. flava*, and *S. pallida* but herbivorous Scaptomyza show reduced sensitivity to some glucosinolates. **(a)** Representative images of scanning electron microscopy (SEM), showing lateral view of the labellum, with illustrations of sensilla type based on length and location. *D. melanogaster* SEM is reprinted from Hiro et al. 2002. **(b)** Left: Schematic of sensilla types present in *D. melanogaster* that may be present in Scaptomyza. Right: Illustration of how single sensilla taste recordings were administered. **(c)** Heatmap of spike responses to caffeine (10mM), lobeline (1mM), sucrose (10mM), across all ∼30 sensilla in each species, illustrating sensilla types by their responses. **(d)** Representatives of each sensilla, class chosen based on hierarchical clustering (**Fig. S2**), were recorded from to generate dose response curves for caffeine, lobeline, and sucrose, and **(e)** for three representative glucosinolates: gluconasturtiin (aromatic), neoglucobrassicin (indolic), and sinigrin (aliphatic). Error bars represent the standard error of the mean. **(f)** Heatplot of spike responses towards all glucosinolates (10mM).

### Identification of sensilla types based on electrophysiology response profiles

To identify herbivore-specific changes in bitter sensitivity in the peripheral nervous system, we first sought to identify whether Scaptomyza species have different functional sensilla classes than *D. melanogaster* (Weiss et al. 2011). In *D. melanogaster* and other *Drosophila* species, sensilla classes can be differentiated based on their electrophysiological responses to caffeine, lobeline, and sucrose (**Fig. 2b**) (Weiss et al. 2011; Dweck et al. 2021; Dweck et al. 2020). Responses to these compounds may have evolved differently in Scaptomyza species, but because all Scaptomyza also show aversion to caffeine and lobeline and attraction to sucrose, similar to *D. melanogaster* (**Fig. 1d**), we reasoned that there may be conservation of functional sensilla classes. Following electrophysiological recordings, we used a hierarchical clustering analysis to assign each of the ∼30 sensilla of each species to a sensilla class.

Responses of all labellar sensilla to caffeine (10 mM), lobeline (1 mM) and sucrose (10 mM) are shown in a heatmap (**Fig. 2c**) and bar plots (**Fig. S1**). L type sensilla across all species responded in a similar manner as those in *D. melanogaster*: unresponsive to bitter compounds (caffeine and lobeline) but strongly stimulated by sugars (sucrose) (**Fig. 2c**). However, the spike responses of L type sensilla were slightly lower in *S. flava* and *S. pallida* compared to *D. melanogaster*. This could be attributed to the fact that *D. melanogaster* feeds mainly on rotting fruits, which are higher in sugars than decaying or living vegetative tissues, which *S. pallida* and *S. flava* feed on, respectively. The I and S type sensilla of *S. flava* and *S. pallida* both responded to bitter compounds and sugars, as they do in *D. melanogaster*. I type sensilla in *D. melanogaster* can be separated into two classes: I-a type, which can be stimulated by lobeline, but not caffeine, and I-b type, which can be stimulated by caffeine but not lobeline (Weiss et al., 2011). In the Scaptomyza species, this distinction between the two types disappeared, although there is some inter-sensilla variation, particularly in *S. pallida* (**Fig. 2c**). The lack of two I types is supported by a hierarchical cluster analysis, which showed almost all I types clustering together in both *S. flava* and *S. pallida* (**Fig. S2**). It is possible that additional bitter compounds could help differentiate I subtypes, but two I subtypes have only been found in *D. melanogaster* and *D. simulans* (Dweck et al. 2020) and no other *Drosophila* (Dweck et al. 2020, Dweck et al. 2021). This pattern suggests that two I types may be derived in the lineage leading to *D. melanogaster* and *D. simulans*. Most notably, while responses to sucrose and lobeline are similar between *S. flava* and *S. pallida* among I types, there is a reduced spike response to caffeine in *S. flava* compared to S. *pallida* (**Fig. 2d**, GLM, t = -2.88, df = 47, P = 0.006), which mirrors their feeding preferences (**Fig. 1d**).

Among S type sensilla, 3 subtypes were present in *S. flava*, S-a, S-b, and S-c (**Fig. 2c**), which are also present in *D. melanogaster*. S-c sensilla have a similar profile as L sensilla as they also do not have a bitter gustatory neuron, and thus only respond to sugars. In *S. flava*, the S-c sensilla corresponded to sensilla S5 and S9 (S4 and S8 in *D. melanogaster*). The same sensilla (S5 and S9) appear to be S-c sensilla in *S. pallida*, and other sensilla may functionally be S-c sensilla as well (S4, S7, S8, S10, S11, S12). In *D. melanogaster*, S-a and S-b sensilla were distinguished by their sensitivity towards caffeine, where the three S-b sensilla are more sensitive to caffeine than the five S-a (Weiss et al. 2011). In *S. flava*, the three S-b sensilla correspond to S4, S7, and S11, that were differentiated from the seven remaining S-a sensilla, not by their sensitivity to caffeine, but by their sensitivity to lobeline, as S-b sensilla exhibited higher spike rates after stimulation with lobeline than S-a (**Fig. 3a**).

**Figure 3.**
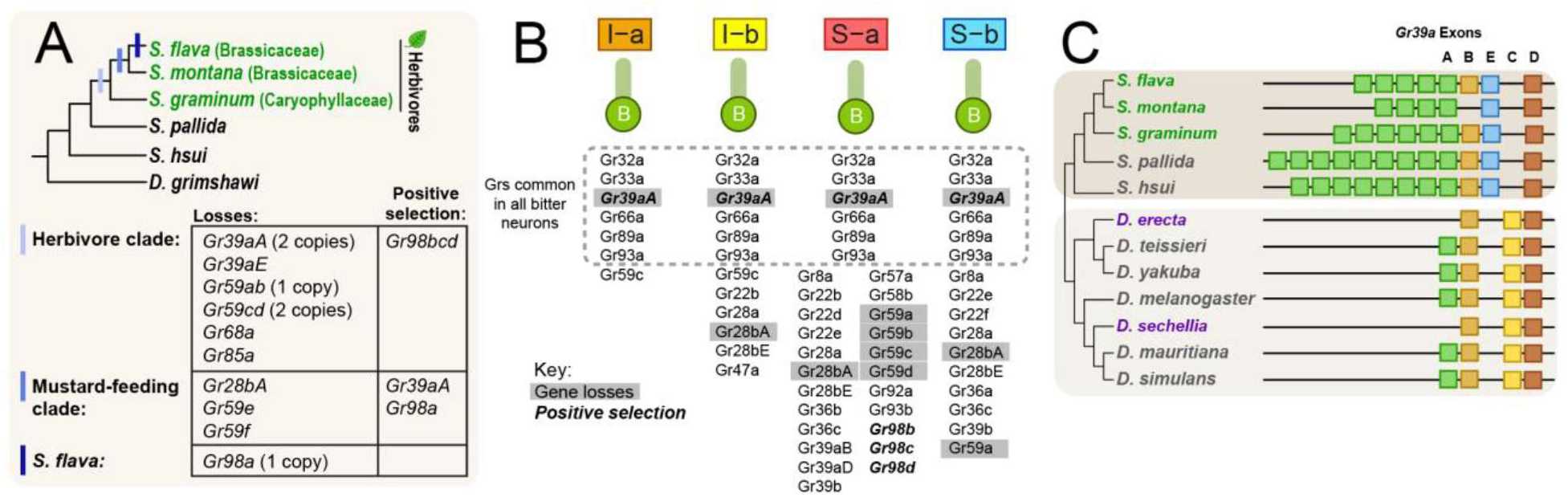
Candidate changes in gustatory receptors in the herbivore *Scaptomyza flava*. **(a)** Genes that have been lost in stepwise progression in the herbivore lineage of Scaptomyza and those that have shown accelerated protein evolution (dN/dS>1 identified from PAML analysis). **(b)** Schematic of bitter gustatory neurons within I and S type sensilla and the expression patterns of homologous GRs from *D. melanogaster*. GRs found in all bitter gustatory neurons in *D. melanogaster* are delineated in the dashed box. GRs that have been lost or pseudogenized in *S. flava*, relative to non-herbivorous Scaptomyza lineages, are highlighted with gray boxes, and those with signatures of relaxed or positive selection are in bold italics. **(c)** Gene structure of Gr39a, showing alternative isoforms A-D, with exon losses of *Gr39aA* occurring across drosophilid herbivores (green text) and other dietary specialists (purple text).

Notably, the sensitivities of both S-a and S-b sensilla towards lobeline in *S. flava* are higher than any sensilla in any of the species we surveyed. This is in congruence with the increased behavioral aversion to lobeline we found in *S. flava* compared to most of the non-herbivores (**Fig. 1d**). In *S. pallida*, based on responses from only caffeine and lobeline, we cannot differentiate S-a and S-b subtypes: all short sensilla that respond to bitter compounds (S1, S2, S3, S6) responded with similar intensity. The delineation of short sensilla types in *S. pallida* is still tentative, and additional compounds are needed to differentiate if any S-b type sensilla exist. Given the conservation of sensilla morphology and spatial locations between *S. flava* and *S. pallida* (**Fig. 2a**), it is possible that the S-b sensilla in *S. pallida* could be the same as those identified in *S. flava*, in which case *S. pallida* S-b sensilla have lost their sensitivity towards bitter compounds.

We chose representative sensilla from each class within each species (**Fig. 2d**) for the remaining recordings. To confirm the sensitivity of each sensilla type towards caffeine, lobeline and sucrose, we generated dose response curves for each representative sensillum. It was not possible to reliably record from *D. melanogaster* S sensilla with 100mM because at high concentrations the spike trains became too noisy for spike identification (Hiroi et al., 2002), so five replicates for these sensilla could not always be achieved. In most cases, spike rates increased with increasing concentrations of the tastant, the only exception being S-c sensilla in *S. flava* and *S. pallida* (**Fig. 2d**). Indeed, we noticed during the initial mapping experiment that there was a small response towards bitter compounds in *S. flava* S-c sensilla (**Fig. 2c**). The negative dose dependent responses, however, suggested that these spikes correspond to the water cells that show higher firing rates in response to lower solute concentrations (Meunier et al., 2003). In *D. melanogaster*, the inclusion of tricholine citrate (>30mM) in the stimulating solution is effective at abolishing water cell firing (Wieczorek & Wolff, 1989). Tricholine citrate was less effective in this regard within the Scaptomyza species. In preliminary recordings, higher concentrations of tricholine citrate were used (up to 300 mM), but there was no reduction in spike rate. Across all sensilla types, spike responses from the water cell were only observed when no other neurons within the sensilla were activated. If sweet or bitter sensitive neurons fired, the spikes from the water cell could be differentiated from these neurons based on the amplitude difference in spikes (**Fig. S3**). These spikes from water cells were not typically seen in I type sensilla, consistent with the lack of water cells in these sensilla (Hiroi et al., 2004).

Lastly, although we only found one I type in Scaptomyza and no S-b type in *S. pallida*, we recorded from sensilla that could represent these sensilla types based on orthologous morphology and location (**Fig. S4a**). The results confirm our initial findings that these sensilla types are no longer present in these species (**Fig. S4b-c**).

### Reduced sensitivity towards mustard-derived glucosinolates

To understand how herbivore responses may have evolved towards host-specific compounds, we performed electrophysiology on the representative sensilla recorded from in **Fig. 2d**. We tested responses towards three glucosinolates, one from each glucosinolate class: gluconasturtiin (aromatic), neoglucobrassicin (indole), and sinigrin (aliphatic). The results were consistent with our findings on their behavioral responses to these compounds. *S. pallida* was strongly sensitive to the glucosinolates gluconasturtiin and neoglucobrassicin, which were both detected by I type and S-a type sensilla (**Fig. 2e**). In contrast, *S. flava* showed marginally reduced sensitivity to these compounds among I type sensilla (**Fig. 2e**; neoglucobrassicin: GLM, t = -2.04, df = 68, P = 0.05; gluconasturtiin: not significant but trending lower). Interestingly, although S-a sensilla detected gluconasturtiin and neoglucobrassicin, there was no reduced sensitivity compared to responses in *S. pallida*, and S-b sensilla showed no response at all to gluconasturtiin and neoglucobrassicin. This lack of response in S-b sensilla in *S. flava* is surprising, given their increased sensitivity towards lobeline (**Fig. 3a** and 3c). These results tenatively suggest that *S. flava*’s loss of aversion to some glucosinolates was achieved through reduced bitter sensitivity in I type sensilla.

*D. melanogaster* showed the lowest levels of aversion in feeding experiments, and their electrophysiological responses towards all three glucosinolates were also extremely low (**Fig. 2e**). Considering *D. melanogaster* did show some glucosinolate aversion, the lack of spike responses from labellar sensilla could indicate that tarsal sensilla play a role in mediating aversion to these compounds. Weak tarsal responses to sinigrin in *D. melanogaster* have been found previously (Ling et al., 2014).

In contrast to aromatic and indolic glucosinolates, the responses to sinigrin, an aliphatic glucosinolate, were significantly different across all sensilla in all species. Both *S. pallida* and *D. melanogaster* showed little sensitivity towards sinigrin across all sensilla. Strikingly, *S. flava* exhibited a positive correlation between sinigrin concentration and spike rate, even in the S-c type sensilla that we would not expect to respond to a bitter compound (**Fig. 2e**). While it is possible that a bitter gustatory neuron could be responding to sinigrin in the labellar sensilla, it may be that a salt-responsive gustatory neuron was stimulated by the salt in the sinigrin solution (Fujishiro et al., 1984). This still begs the question as to how species like *S. pallida*, *D. melanogaster*, and *S. flava* detect glucosinolates like sinigrin. As mentioned earlier, in *D. melanogaster*, singrin is detected by tarsal sensilla (Ling et al., 2014), and it could be possible that some glucosinolates, like sinigrin, are only detected by tarsi. An alternative hypothesis is that sinigrin aversion may be caused by direct or indirect effects of sinigrin on sweet neuron spiking. We addressed this with recordings involving solutions filled with both sucrose and sinigrin, and tested high (10mM) and low (1mM) sinigrin concentrations. There were significant differences between the spike rate between high and low sinigrin concentrations in all three species (**Fig. 4e**; *D. melanogaster*: t(19) = 4.15, P < 0.001; *S. pallida*: t(10) = 4.47, P = 0.001; *S. flava*: t(15) = 2.11, P = 0.05), which suggests that sinigrin (and potentially other glucosinolates) may be inhibiting sweet neuron firing as well. While this is one mechanism in which aliphatic glucosinolates may elicit aversion, it remains unclear whether they are directly detected by gustatory receptors.

We performed electrophysiological recordings on three additional glucosinolates, one from each glucosinolate class - glucotropaeolin (aromatic), glucobrassicin (indolic), and glucoraphanin (aliphatic) - to test whether I and S-a responses were consistent within glucosinolate classes. Spike rates varied among the aromatic and indolic glucosinolates (all tested at 10mM), but *S. flava* had marginally lower spike rates than *S. pallida* among I sensilla (**Fig. 2f** and **Fig. S4d**). Similarly, responses to the aliphatic glucosinolate glucoraphanin were low, as with sinigrin.

### Candidate gustatory receptor genetic changes

We next sought to identify candidate genetic changes in gustatory receptors associated with the reduced bitter sensitivity found in *S. flava*. We examined copy number changes (i.e. duplications, losses or pseudogenizations) and tested for evidence of positive selection tested through a maximum likelihood approach using PAML. In these analyses, we focused on changes along the basal branch preceding all herbivore lineages (*S. flava*, *S. montana*, and *S. graminum*), along the branch leading to both species that feed on mustard plants (*S. flava* and *S. montana*), and to *S. flava* only.

As reported in our previously published study, we identified several GRs lost among all of the herbivorous Scaptomyza (two *Gr39aA*-specific exons, the *Gr39aE*-specific exon, one paralog of *Gr59ab*, two paralogs of *Gr59cd, Gr68a*, and *Gr85a*), three GRs lost among the mustard feeders (*Gr28bA, Gr59e*, and *Gr59f*), and one GR along the lineage to *S. flava* (one paralog of Gr98a) (**Fig. 3a**).

We used a codon-based branch-site test for positive selection, and identified genes with elevated dN/dS values at a proportion of sites along one of the herbivore branches described above. We found evidence for positive selection within *Gr98bcd* along the branches preceding all three herbivores (ω_2b_ = 179.55), and along the branch leading to mustard feeders within the *Gr39aA*-specific exons (ω_2b_ = 13.71), and within copies of Gr98a (ω_2b_ = 4.83) (**Fig. 3a**; Table 2). The results of all branch-site tests are present in Table S7.

**Table 2.** Maximum likelihood tests for positive selection

| Gene | Foreground branch | dN/dS (proportion of sites) |  |  |  | lnL | LRT (2*ΔlnL) | q-value (5% FDR) |
| --- | --- | --- | --- | --- | --- | --- | --- | --- |
| Gr98bcd | herbivore | w0= | 0.16 (73%) | w1= | 1 (27%) | -12238.2 | 10.2 | 0.046 |
|  |  | w2a= | 179.55 (0.4%) | w2b = | 179.55 (0.1%) |  |  |  |
| Gr39aA | mustard | w0= | 0.13 (50%) | w1= | 1 (49%) | -31858.5 | 30.8 | 8.70E-06 |
|  |  | w2a= | 13.71 (0.2%) | w2b = | 13.71 (0.2%) |  |  |  |
| Gr98a | mustard | w0= | 0.22 (69%) | w1= | 1 (23%) | -19590 | 11.2 | 0.035 |
|  |  | w2a= | 4.83 (6%) | w2b<br>= | 4.83 (2%) |  |  |  |

## DISCUSSION

Understanding how herbivore specialists evolve feeding preferences for their new host plants, particularly towards host plant defense compounds, is key to understanding their adaptation to this new niche. Here, we investigated how the taste preferences of a mustard specialist, *S. flava*, have evolved towards glucosinolates, compounds specific to the flowering plant order Brassicales. We found that *S. flava* is less averse to glucosinolates, and this appears to be attributable to the reduced sensitivity of bitter neurons within the I type sensilla towards glucosinolates and other bitter compounds (i.e., caffeine).

While most generalist herbivores and non-herbivores find glucosinolates aversive, many mustard specialists have evolved attraction towards these compounds. We confirmed that non-herbivorous drosophilids (*S. pallida*, *S. hsui*, and *D. melanogaster*) are averse to glucosinolates, with *S. pallida* and S. shui showing especially strong sensitivity to glucosinolates. These findings are consistent with the fact that these two Scaptomyza non-herbivores likely encounter these compounds while feeding on decaying plant material. However, we also found that the young mustard specialist *S. flava* was still averse to higher concentrations of glucosinolates, and this aversion, in Scaptomyza herbivores and non-herbivores alike, is mediated by specific sensilla classes that house bitter gustatory neurons, specifically I type and S-a type sensilla. Maintaining the capability of sensing glucosinolates is not surprising given that *S. flava* is still an evolutionarily young herbivore lineage. It is still directly influenced by the toxic effects of these host plant defenses, showing slowed development and reduced weight gain from feeding on glucosinolates (Gloss et al., 2019). Moreover, *S. flava* is unique among studied Brassicales specialists, like cabbage white butterflies and diamondback moths, in not having a physiological mechanism that prevents the formation of toxic isothiocyanates and other hydrolysis products in its body. Drosophilids, including *S. flava*, *D. melanogaster* and humans, use the ancient mercapturic acid pathway to detoxify isothiocyanates (Gloss et al., 2019). At lower glucosinolate concentrations, *S. flava* has lost or reduced its aversion to most of the glucosinolates tested, which enables these herbivores to feed on plants harboring lower levels of these compounds. Estimates of glucosinolate concentrations from leaves have been reported in the range of 0-10mM in some plant species (Merritt, 1996). But glucosinolate concentrations can vary depending on the age of the plant, the species, the tissue type, and environmental conditions (Brown et al., 2003; Feeny & Rosenberry, 1982; Wentzell & Kliebenstein, 2008). The reduced glucosinolate sensitivity in *S. flava* would thus allow them to sense and avoid higher, toxic concentrations, while readily feeding on lower, less toxic concentrations.

A lingering question is whether there is gustatory attraction towards any glucosinolates - potentially in a context-dependent manner (e.g., positional attraction or oviposition stimulation) in S. *flava.* We know from *D. melanogaster* that the same compounds that can be aversive to feeding can also be stimulatory for egg-laying through differential expression in different gustatory tissues (Joseph & Heberlein, 2012). While we only found evidence for no preference or slight aversion to the tested glucosinolates in *S. flava*, previous studies assessing the choices of *S. flava* across 585 genotyped accessions of *Arabidopsis thaliana* indicate that aliphatic and indolic glucosinolates can positively influence whether and how much they feed on a given plant (Gloss et al. 2017). Gustatory synergism caused by a blend of plant-derived compounds may influence *S. flava* feed and/or oviposit, as seen in other insect species (Hojo et al. 2008). Volatile isothiocyanates activate olfactory receptors in *S. flava* that are specifically tuned to these mustard oils - mediated by recently triplicated copies of the odorant receptor gene *Or67b* (Matsunaga et al. 2021). Further study will be needed to test whether olfactory cues alone stimulate feeding and egg-laying in herbivorous Scaptomyza, or whether additional gustatory cues are necessary to elicit these behaviors.

We suspected that there would be differences in the responses of mustard feeders towards different classes of glucosinolates (aliphatic, aromatic, and indolic). Over 120 glucosinolates have been found naturally occurring in mustard plants, and this chemical diversity has evolved to evade herbivore counter-adaptations (Fahey et al., 2001). We expected that mustard specialists may evolve greater losses of aversion towards indolic glucosinolates because the isothiocyanates they form are unstable and break down rapidly (Agerbirk et al., 1998; Chevolleau et al., 1997), leaving potentially less toxic end products like indole-3-carbinol. We found feeding and physiological responses varied across different glucosinolates, but testing with more compounds and at more concentrations is needed to conclusively determine whether class differences exist.

The diversity of plant secondary metabolites is matched by the diversity of GRs that have evolved in insects to detect bitter compounds: out of the roughly 68 GR proteins in *D. melanogaster*, seven of them function to detect sugars (Dahanukar et al., 2007; Fujii et al., 2015; Slone et al., 2007), while almost half of the remaining GRs are expressed in bitter gustatory neurons, with many playing a functional role in detecting bitter compounds (Lee et al., 2009, 2012; Moon et al., 2006; Sung et al., 2017; Weiss et al., 2011). While we still do not know which GRs are expressed in each gustatory neuron within the labellar sensilla in *S. flava*, we can leverage the homology with GR expression among the bitter gustatory neurons of *D. melanogaster*. Despite the limitations that GR expression may have evolved in Scaptomyza from its ancestor with *D. melanogaster*, including the presence of only a single I type of sensilla, we are still able to highlight several candidate genetic changes in key GRs that may underpin evolved behavioral adaptations in glucosinolate detection.

Our single sensillum recordings showed a reduction in sensitivity towards some glucosinolates and caffeine among I type sensilla. Among the GR losses and GRs for which there is evidence of positive selection acting on them, *Gr39aA* and *Gr28bA* are the only GRs expressed in the I type sensilla of *D. melanogaster* (as well as in S-type) (**Fig. 3b**). In *D. melanogaster*, *Gr28bA* has been implicated in the detection of saponin, which are triterpene glycosides found in a variety of flowering plant families (Sang et al., 2019). Thus, its loss from mustard-feeding Scaptomyza may drive some loss of bitter detection. *Gr39aA* is likely an even stronger candidate. An alternative isoform of Gr39a, it is critically important for the detection of a wide variety of bitter compounds and is found in all bitter gustatory neurons (H. K. M. Dweck & Carlson, 2020; Weiss et al., 2011). Only a single copy of the exon that encodes *Gr39aA* is found in *D. melanogaster*, but unlike the other four exons that encode alternative isoforms, the A exon has undergone numerous duplications in many *Drosophila* species (Gardiner et al., 2008), including *S. pallida* and *S. hsui* which have nine and eight copies each, respectively (**Fig. 3c**). *S. flava* has six, and *S. montana*, an even more specialized mustard feeder than *S. flava*, has only four copies. Based on the loss of *Gr39aA* exons and positive selection on the remaining copies, the number of *Gr39aA* copies could impact regulatory expression of this GR. The role of this GR in spurring changes in bitter detection is further suggested by losses of *Gr39aA* in several other specialist lineages, including in the noni specialist *D. sechellia* and Pandanus spp. fruit specialists *D. erecta*. Interestingly, the loss of Gr39a in *D. melanogaster* caused a reduction in caffeine sensitivity in I-b type sensilla, while also causing an increase in lobeline sensitivity in I-a type sensilla (Dweck et al. 2020), a pattern similarly seen in our single sensillum recordings (**Fig. 2d**). Increased sensitivity due to GR loss has been shown for several other GRs and compounds as well (H. K. M. Dweck & Carlson, 2020), although the mechanisms causing this are still unknown.

In summary, our study supports an evolutionary model wherein reduced bitter sensitivity at the level of sensory neurons has enabled an herbivorous drosophild to feed on living vegetative plants tissues rife with bitter and toxic plant defense chemicals. The redundancy of the gustatory taste system has allowed these flies to maintain some bitter sensitivity, critical for insects still inhabiting complex environments in which they come into contact with diverse toxins. With the identification of candidate GRs that have been lost in *S. flava*, particularly *Gr39aA*, it will be interesting to functionally test the role of these genetic changes in driving feeding preferences.

## ACKNOWLEDGMENTS

We thank Susan Bernstein for assistance in rearing *Arabidopsis* plants; Peter Oboyski, Pénélope Tarapacki, and Béatrice Denis for their assistance in growing *S. flava* and *Arabidopsis*; Kristin Scott, Peter Sudmant, Doris Bachtrog, Carolina Reisenman, Teruyuki Matsunaga, and other members of the Whiteman Lab for their feedback. We also acknowledge Jacobs Hall for utilizing their resources for 3D printing parts of the fly chambers for feeding experiments.

