## Supplemental Tables & Figures for "Taste evolution in an herbivorous drosophilid"

**Supplementary Material for Evolution of gustatory changes in herbivorous *Drosophila***

### 1. SUPPLEMENTARY TABLES

**Table S1.** Summary statistics of feeding preferences towards caffeine and lobeline (**Fig. 1d**).

| | | Summary statistics | | | | | | Kruskal-Wallis test | | | Post-hoc Dunn's test | | | | | |
|---|---|---|---|---|---|---|---|---|---|---|---|---|---|---|---|---|
| | **Conc. (mM)** | **Species** | **N** | **Preference Index** | **sd** | **se** | **ci** | **chi-squared** | **df** | | **Compari-son** | **Z** | **P.unadj** | **P.adj** | **Group** | **Letter** |
| Caffeine | 1 | Dmel | 12 | 0.541 | 0.144 | 0.041 | 0.091 | 23.026 | 3 | **3.99E-05** | Dmel-Sfla | 1.200 | 0.23 | 1.000 | Dmel | a |
| | | Shsu | 9 | 0.186 | 0.19 | 0.063 | 0.146 | | | | Dmel-Shsu | 3.583 | 0.0003 | **0.002** | Sfla | ab |
| | | Spal | 11 | 0.138 | 0.108 | 0.033 | 0.073 | | | | Sfla-Shsu | 1.860 | 0.063 | 0.378 | Shsu | bc |
| | | Sfla | 6 | 0.396 | 0.102 | 0.042 | 0.107 | | | | Dmel-Spal | 4.312 | 0.00002 | **0.0001** | Spal | c |
| | | | | | | | | | | | Sfla-Spal | 2.365 | 0.018 | 0.108 | | |
| | | | | | | | | | | | Shsu-Spal | 0.489 | 0.624 | 1.000 | | |
| | 10 | Dmel | 14 | 0.083 | 0.136 | 0.036 | 0.078 | 1.566 | 3 | 0.667 | Dmel-Sfla | 0.463 | 0.643 | 1.000 | Dmel | a |
| | | Shsu | 12 | 0.142 | 0.195 | 0.056 | 0.124 | | | | Dmel-Shsu | -0.419 | 0.675 | 1.000 | Sfla | a |
| | | Spal | 11 | 0.132 | 0.166 | 0.05 | 0.112 | | | | Sfla-Shsu | -0.763 | 0.445 | 1.000 | Shsu | a |
| | | Sfla | 5 | 0.052 | 0.097 | 0.043 | 0.12 | | | | Dmel-Spal | -0.919 | 0.358 | 1.000 | Spal | a |
| | | | | | | | | | | | Sfla-Spal | -1.134 | 0.257 | 1.000 | | |
| | | | | | | | | | | | Shsu-Spal | -0.492 | 0.623 | 1.000 | | |
| Lobeline | 0.1 | Dmel | 7 | 0.182 | 0.057 | 0.022 | 0.053 | 13.071 | 3 | **0.004** | Dmel-Sfla | 2.596 | 0.009 | 0.057 | Dmel | a |
| | | Shsu | 7 | 0.047 | 0.098 | 0.037 | 0.091 | | | | Dmel-Shsu | 2.362 | 0.018 | 0.109 | Sfla | b |
| | | Spal | 9 | 0.216 | 0.185 | 0.062 | 0.142 | | | | Sfla-Shsu | -0.440 | 0.66 | 1.000 | Shsu | b |
| | | Sfla | 5 | 0.013 | 0.03 | 0.013 | 0.037 | | | | Dmel-Spal | -0.002 | 0.998 | 1.000 | Spal | a |
| | | | | | | | | | | | Sfla-Spal | -2.727 | 0.006 | **0.038** | | |
| | | | | | | | | | | | Shsu-Spal | -2.507 | 0.012 | 0.073 | | |
| | 1 | Dmel | 12 | 0.015 | 0.038 | 0.011 | 0.024 | 7.819 | 3 | 0.05 | Dmel-Sfla | 0.867 | 0.386 | 1.000 | Dmel | ab |
| | | Shsu | 5 | 0 | 0 | 0 | 0 | | | | Dmel-Shsu | 0.814 | 0.415 | 1.000 | Sfla | a |
| | | Spal | 7 | 0.044 | 0.075 | 0.028 | 0.07 | | | | Sfla-Shsu | 0.000 | 1 | 1.000 | Shsu | a |
| | | Sfla | 6 | 0 | 0 | 0 | 0 | | | | Dmel-Spal | -1.921 | 0.055 | 0.329 | Spal | b |
| | | | | | | | | | | | Sfla-Spal | -2.421 | 0.015 | 0.093 | | |
| | | | | | | | | | | | Shsu-Spal | -2.300 | 0.021 | 0.129 | | |

**Table S2.** Summary statistics of feeding preferences towards glucosinolates (**Fig. 1e**).

| | | Summary statistics | | | | | | Kruskal-Wallis test | | | Post-hoc Dunn's test | | | | | |
|---|---|---|---|---|---|---|---|---|---|---|---|---|---|---|---|---|
| **Compound** | **Conc. (mM)** | **Species** | **N** | **Preference Index** | **sd** | **se** | **ci** | **chi-squared** | **df** | **p-figure** | **Comparison** | **Z** | **P.unadj** | **P.adj** | **Group** | **Letter** |
| Gluconasturtiin | 1 | Dmel | 21 | 0.327 | 0.199 | 0.043 | 0.09 | 9.839 | 3 | **0.02** | Dmel-Sfla | -2.89 | 0.004 | **0.023** | Dmel | a |
| | | Shsu | 12 | 0.345 | 0.179 | 0.052 | 0.114 | | | | Dmel-Shsu | -0.289 | 0.773 | 1 | Sfla | b |
| | | Spal | 18 | 0.312 | 0.202 | 0.048 | 0.1 | | | | Sfla-Shsu | 2.506 | 0.012 | 0.073 | Shsu | a |
| | | Sfla | 5 | 0.662 | 0.084 | 0.038 | 0.105 | | | | Dmel-Spal | 0.301 | 0.764 | 1 | Spal | a |
| | | | | | | | | | | | Sfla-Spal | 3.036 | 0.002 | **0.014** | | |
| | | | | | | | | | | | Shsu-Spal | 0.539 | 0.59 | 1 | | |
| | 10 | Dmel | 10 | 0.297 | 0.187 | 0.059 | 0.134 | 17.405 | 3 | **0.001** | Dmel-Sfla | 0.819 | 0.413 | 1 | Dmel | a |
| | | Shsu | 9 | 0.02 | 0.026 | 0.009 | 0.02 | | | | Dmel-Shsu | 3.972 | 0.00007 | **0.0004** | Sfla | a |
| | | Spal | 8 | 0.176 | 0.168 | 0.059 | 0.14 | | | | Sfla-Shsu | 3.174 | 0.002 | **0.009** | Shsu | b |
| | | Sfla | 10 | 0.189 | 0.126 | 0.04 | 0.09 | | | | Dmel-Spal | 1.398 | 0.162 | 0.973 | Spal | a |
| | | | | | | | | | | | Sfla-Spal | 0.626 | 0.532 | 1 | | |
| | | | | | | | | | | | Shsu-Spal | -2.391 | 0.017 | 0.101 | | |
| Glucotropaeolin | 1 | Dmel | 10 | 0.554 | 0.121 | 0.038 | 0.086 | 13.529 | 3 | **0.004** | Dmel-Sfla | 1.338 | 0.181 | 1 | Dmel | a |
| | | Shsu | 7 | 0.368 | 0.132 | 0.05 | 0.122 | | | | Dmel-Shsu | 2.021 | 0.043 | 0.26 | Sfla | ab |
| | | Spal | 6 | 0.244 | 0.078 | 0.032 | 0.082 | | | | Sfla-Shsu | 0.548 | 0.584 | 1 | Shsu | bc |
| | | Sfla | 6 | 0.434 | 0.175 | 0.071 | 0.183 | | | | Dmel-Spal | 3.613 | 0.0003 | **0.002** | Spal | c |
| | | | | | | | | | | | Sfla-Spal | 2.035 | 0.042 | 0.251 | | |
| | | | | | | | | | | | Shsu-Spal | 1.564 | 0.118 | 0.708 | | |
| | 10 | Dmel | 8 | 0.247 | 0.107 | 0.038 | 0.089 | 13.069 | 3 | **0.004** | Dmel-Sfla | 0.535 | 0.593 | 1 | Dmel | a |
| | | Shsu | 5 | 0.104 | 0.133 | 0.059 | 0.165 | | | | Dmel-Shsu | 1.926 | 0.054 | 0.324 | Sfla | a |
| | | Spal | 7 | 0.029 | 0.038 | 0.014 | 0.035 | | | | Sfla-Shsu | 1.457 | 0.145 | 0.87 | Shsu | ab |
| | | Sfla | 8 | 0.22 | 0.185 | 0.066 | 0.155 | | | | Dmel-Spal | 3.265 | 0.001 | **0.007** | Spal | b |
| | | | | | | | | | | | Sfla-Spal | 2.749 | 0.006 | **0.036** | | |
| | | | | | | | | | | | Shsu-Spal | 1.011 | 0.312 | 1 | | |
| Neoglucobrassicin | 1 | Dmel | 9 | 0.584 | 0.189 | 0.063 | 0.145 | 6.972 | 3 | 0.073 | Dmel-Sfla | 0.756 | 0.45 | 1 | Dmel | a |
| | | Shsu | 7 | 0.371 | 0.199 | 0.075 | 0.184 | | | | Dmel-Shsu | 1.947 | 0.051 | 0.309 | Sfla | ab |
| | | Spal | 4 | 0.3 | 0.127 | 0.063 | 0.201 | | | | Sfla-Shsu | 0.172 | 0.863 | 1 | Shsu | ab |
| | | Sfla | 1 | 0.4 | NA | NA | NaN | | | | Dmel-Spal | 2.366 | 0.018 | 0.108 | Spal | b |
| | | | | | | | | | | | Sfla-Spal | 0.559 | 0.576 | 1 | | |
| | | | | | | | | | | | Shsu-Spal | 0.703 | 0.482 | 1 | | |
| | 10 | Dmel | 8 | 0.218 | 0.179 | 0.063 | 0.15 | 11.862 | 3 | **0.008** | Dmel-Sfla | -1.329 | 0.184 | 1 | Dmel | a |
| | | Shsu | 7 | 0.118 | 0.093 | 0.035 | 0.086 | | | | Dmel-Shsu | 0.843 | 0.399 | 1 | Sfla | a |
| | | Spal | 6 | 0.022 | 0.036 | 0.015 | 0.038 | | | | Sfla-Shsu | 1.937 | 0.053 | 0.317 | Shsu | ab |
| | | Sfla | 3 | 0.398 | 0.037 | 0.021 | 0.092 | | | | Dmel-Spal | 2.522 | 0.012 | 0.07 | Spal | b |
| | | | | | | | | | | | Sfla-Spal | 3.199 | 0.001 | **0.008** | | |
| | | | | | | | | | | | Shsu-Spal | 1.663 | 0.096 | 0.578 | | |

| | | | | | | | | | | | | | | | | |
|---|---|---|---|---|---|---|---|---|---|---|---|---|---|---|---|---|
| Glucobrassicin | 1 | Dmel | 10 | 0.462 | 0.207 | 0.065 | 0.148 | 12.388 | 3 | **0.006** | Dmel-Sfla | 0.174 | 0.862 | 1 | Dmel | a |
| | | Shsu | 6 | 0.809 | 0.11 | 0.045 | 0.115 | | | | Dmel-Shsu | -2.948 | 0.003 | **0.019** | Sfla | a |
| | | Spal | 6 | 0.414 | 0.196 | 0.08 | 0.206 | | | | Sfla-Shsu | -2.971 | 0.003 | **0.018** | Shsu | b |
| | | Sfla | 8 | 0.403 | 0.265 | 0.094 | 0.222 | | | | Dmel-Spal | 0.37 | 0.711 | 1 | Spal | a |
| | | | | | | | | | | | Sfla-Spal | 0.202 | 0.84 | 1 | | |
| | | | | | | | | | | | Shsu-Spal | 2.968 | 0.003 | **0.018** | | |
| | 10 | Dmel | 11 | 0.158 | 0.16 | 0.048 | 0.108 | 9.493 | 3 | **0.023** | Dmel-Sfla | -2.052 | 0.04 | 0.241 | Dmel | a |
| | | Shsu | 8 | 0.312 | 0.315 | 0.111 | 0.263 | | | | Dmel-Shsu | -0.544 | 0.586 | 1 | Sfla | b |
| | | Spal | 11 | 0.075 | 0.095 | 0.029 | 0.064 | | | | Sfla-Shsu | 1.429 | 0.153 | 0.918 | Shsu | ab |
| | | Sfla | 7 | 0.484 | 0.318 | 0.12 | 0.294 | | | | Dmel-Spal | 1.111 | 0.266 | 1 | Spal | a |
| | | | | | | | | | | | Sfla-Spal | 3.032 | 0.002 | **0.015** | | |
| | | | | | | | | | | | Shsu-Spal | 1.564 | 0.118 | 0.707 | | |
| Sinigrin | 1 | Dmel | 15 | 0.459 | 0.135 | 0.035 | 0.075 | 4.756 | 3 | 0.191 | Dmel-Sfla | -0.584 | 0.56 | 1 | Dmel | a |
| | | Shsu | 11 | 0.381 | 0.218 | 0.066 | 0.147 | | | | Dmel-Shsu | 1.209 | 0.227 | 1 | Sfla | a |
| | | Spal | 13 | 0.373 | 0.164 | 0.045 | 0.099 | | | | Sfla-Shsu | 1.501 | 0.133 | 0.8 | Shsu | a |
| | | Sfla | 6 | 0.513 | 0.193 | 0.079 | 0.202 | | | | Dmel-Spal | 1.568 | 0.117 | 0.701 | Spal | a |
| | | | | | | | | | | | Sfla-Spal | 1.775 | 0.076 | 0.455 | | |
| | | | | | | | | | | | Shsu-Spal | 0.279 | 0.78 | 1 | | |
| | 10 | Dmel | 28 | 0.455 | 0.134 | 0.025 | 0.052 | 37.447 | 3 | **0** | Dmel-Sfla | 4.09 | 0.00004 | **0.0003** | Dmel | a |
| | | | | | | | | | | | | | 0.000000 | | | |
| | | Shsu | 20 | 0.114 | 0.155 | 0.035 | 0.073 | | | | Dmel-Shsu | 5.39 | 07 | **0.0000004** | Sfla | b |
| | | Spal | 24 | 0.247 | 0.19 | 0.039 | 0.08 | | | | Sfla-Shsu | -0.815 | 0.415 | 1 | Shsu | b |
| | | Sfla | 5 | 0.044 | 0.099 | 0.044 | 0.123 | | | | Dmel-Spal | 3.376 | 0.0007 | **0.004** | Spal | c |
| | | | | | | | | | | | Sfla-Spal | -2.129 | 0.033 | 0.2 | | |
| | | | | | | | | | | | Shsu-Spal | -2.11 | 0.035 | 0.209 | | |
| Glucoraphanin | 1 | Dmel | 12 | 0.541 | 0.216 | 0.062 | 0.137 | 4.844 | 3 | 0.184 | Dmel-Sfla | 1.718 | 0.086 | 0.515 | Dmel | a |
| | | Shsu | 9 | 0.425 | 0.239 | 0.08 | 0.184 | | | | Dmel-Shsu | 1.41 | 0.159 | 0.951 | Sfla | ab |
| | | Spal | 12 | 0.349 | 0.147 | 0.042 | 0.093 | | | | Sfla-Shsu | -0.288 | 0.773 | 1 | Shsu | ab |
| | | Sfla | 9 | 0.375 | 0.22 | 0.073 | 0.169 | | | | Dmel-Spal | 1.981 | 0.048 | 0.286 | Spal | b |
| | | | | | | | | | | | Sfla-Spal | 0.116 | 0.908 | 1 | | |
| | | | | | | | | | | | Shsu-Spal | 0.424 | 0.672 | 1 | | |
| | 10 | Dmel | 8 | 0.442 | 0.211 | 0.075 | 0.177 | 6.89 | 3 | 0.075 | Dmel-Sfla | 2.217 | 0.027 | 0.16 | Dmel | a |
| | | Shsu | 8 | 0.225 | 0.132 | 0.047 | 0.11 | | | | Dmel-Shsu | 2.234 | 0.026 | 0.153 | Sfla | b |
| | | Spal | 8 | 0.292 | 0.172 | 0.061 | 0.144 | | | | Sfla-Shsu | -0.258 | 0.797 | 1 | Shsu | b |
| | | Sfla | 5 | 0.2 | 0.158 | 0.071 | 0.197 | | | | Dmel-Spal | 1.514 | 0.13 | 0.781 | Spal | ab |
| | | | | | | | | | | | Sfla-Spal | -0.889 | 0.374 | 1 | | |
| | | | | | | | | | | | Shsu-Spal | -0.72 | 0.471 | 1 | | |

**Table S3.** Summary statistics of electrophysiological responses to sucrose, caffeine, and lobeline using generalized linear models with gamma distributions **(Fig. 2d).** Spikes per second from labellar sensilla was regressed against species, concentration, and their interaction term.

| Compound | Sensilla | | Estimate | Std. Error | t value | Pr(>\|t\|) | | Sensilla | | Estimate | Std. Error | t value | Pr(>\|t\|) | |
|---|---|---|---|---|---|---|---|---|---|---|---|---|---|---|
| caffeine | L | (Intercept) | 0.619 | 0.095 | 6.549 | **<0.001** | *** | S-a | (Intercept) | 0.064 | 0.011 | 5.655 | **<0.001** | *** |
| | | insectDmel | 0.246 | 0.157 | 1.571 | 0.123 | | | insectDmel | 0.03 | 0.021 | 1.421 | 0.16 | |
| | | insectSpal | 0.281 | 0.173 | 1.624 | 0.112 | | | insectSpal | 0.029 | 0.019 | 1.522 | 0.133 | |
| | | conc1 | 0.001 | 0.002 | 0.564 | 0.575 | | | conc1 | -0.0004 | <0.001 | -3.237 | **0.002** | ** |
| | | insectDmel:conc1 | -0.001 | 0.003 | -0.434 | 0.666 | | | insectDmel:conc1 | 0.0001 | <0.001 | 0.32 | 0.75 | |
| | | insectSpal:conc1 | <0.001 | 0.003 | 0.036 | 0.971 | | | insectSpal:conc1 | -0.0002 | <0.001 | -0.937 | 0.352 | |
| | I-a | (Intercept) | 0.148 | 0.031 | 4.691 | **<0.001** | *** | S-b | (Intercept) | 0.046 | 0.008 | 5.582 | **<0.001** | *** |
| | | insectDmel | 0.268 | 0.111 | 2.419 | **0.019** | * | | insectDmel | 0.005 | 0.013 | 0.38 | 0.705 | |
| | | insectSpal | -0.095 | 0.033 | -2.879 | **0.006** | ** | | insectSpal | 0.101 | 0.031 | 3.279 | **0.002** | ** |
| | | conc1 | -0.001 | <0.001 | -3.301 | **0.002** | ** | | conc1 | -0.0002 | <0.001 | -1.32 | 0.192 | |
| | | insectDmel:conc1 | 0.004 | 0.003 | 1.549 | 0.127 | | | insectDmel:conc1 | 0.000003 | <0.001 | 0.018 | 0.986 | |
| | | insectSpal:conc1 | 0.001 | <0.001 | 2.567 | **0.013** | * | | insectSpal:conc1 | 0.002 | 0.001 | 1.458 | 0.15 | |
| | I-b | (Intercept) | 0.11 | 0.023 | 4.834 | **<0.001** | *** | S-c | (Intercept) | 0.064 | 0.022 | 2.885 | **0.006** | ** |
| | | insectDmel | -0.065 | 0.024 | -2.687 | **0.01** | ** | | insectDmel | 0.701 | 0.216 | 3.249 | **0.002** | ** |
| | | insectSpal | -0.057 | 0.025 | -2.312 | **0.025** | * | | insectSpal | 0.071 | 0.044 | 1.608 | 0.114 | |
| | | conc1 | -0.001 | <0.001 | -2.341 | **0.024** | * | | conc1 | 0.001 | 0.001 | 1.34 | 0.186 | |
| | | insectDmel:conc1 | 0.001 | 0 | 2.047 | **0.046** | * | | insectDmel:conc1 | 0.001 | 0.005 | 0.242 | 0.81 | |
| | | insectSpal:conc1 | <0.001 | <0.001 | 1.429 | 0.16 | | | insectSpal:conc1 | 0.001 | 0.002 | 0.752 | 0.456 | |
| lobeline | L | (Intercept) | 0.826 | 0.11 | 7.525 | **<0.001** | *** | S-a | (Intercept) | 0.03 | 0.004 | 8.392 | **<0.001** | *** |
| | | insectDmel | 0.174 | 0.158 | 1.107 | 0.274 | | | insectDmel | 0.01 | 0.006 | 1.573 | 0.121 | |
| | | insectSpal | -0.034 | 0.148 | -0.232 | 0.818 | | | insectSpal | 0.029 | 0.007 | 3.959 | **<0.001** | *** |
| | | conc1 | 0.002 | 0.019 | 0.109 | 0.913 | | | conc1 | -0.001 | 0.001 | -2.467 | **0.017** | * |
| | | insectDmel:conc1 | -0.002 | 0.027 | -0.077 | 0.939 | | | insectDmel:conc1 | 0.002 | 0.001 | 2.357 | **0.022** | * |
| | | insectSpal:conc1 | 0.005 | 0.025 | 0.18 | 0.858 | | | insectSpal:conc1 | -0.001 | 0.001 | -1.428 | 0.158 | |
| | I-a | (Intercept) | 0.056 | 0.008 | 6.948 | **<0.001** | *** | S-b | (Intercept) | 0.021 | 0.005 | 4.245 | **<0.001** | *** |
| | | insectDmel | 0.022 | 0.015 | 1.488 | 0.142 | | | insectDmel | 0.016 | 0.009 | 1.703 | 0.094 | . |
| | | insectSpal | 0.002 | 0.012 | 0.171 | 0.865 | | | insectSpal | 0.106 | 0.029 | 3.728 | **<0.001** | *** |
| | | conc1 | -0.002 | 0.001 | -1.775 | 0.081 | . | | conc1 | -0.0005 | 0.001 | -0.645 | 0.521 | |
| | | insectDmel:conc1 | 0.006 | 0.003 | 2.364 | **0.021** | * | | insectDmel:conc1 | 0.0001 | 0.002 | 0.065 | 0.949 | |
| | | insectSpal:conc1 | 0.001 | 0.002 | 0.375 | 0.709 | | | insectSpal:conc1 | -0.001 | 0.005 | -0.19 | 0.85 | |
| | I-b | (Intercept) | 0.044 | 0.009 | 4.99 | **<0.001** | *** | S-c | (Intercept) | 0.087 | 0.03 | 2.933 | **0.005** | ** |
| | | insectDmel | 0.433 | 0.09 | 4.796 | **<0.001** | *** | | insectDmel | 0.529 | 0.197 | 2.683 | **0.01** | ** |
| | | insectSpal | 0.011 | 0.013 | 0.794 | 0.431 | | | insectSpal | 0.099 | 0.071 | 1.399 | 0.168 | |
| | | conc1 | -0.001 | 0.001 | -0.509 | 0.613 | | | conc1 | 0.019 | 0.011 | 1.722 | 0.091 | . |
| | | insectDmel:conc1 | 0.047 | 0.026 | 1.796 | 0.078 | . | | insectDmel:conc1 | -0.01 | 0.034 | -0.291 | 0.772 | |
| | | insectSpal:conc1 | -0.001 | 0.002 | -0.739 | 0.463 | | | insectSpal:conc1 | 0.055 | 0.042 | 1.299 | 0.2 | |

| | | | | | | | | | | | | | | |
|---|---|---|---|---|---|---|---|---|---|---|---|---|---|---|
| sucrose | L | (Intercept) | 0.0365 | 0.007 | 5.252 | **<0.001** | *** | S-a | (Intercept) | 0.06 | 0.011 | 5.624 | **<0.001** | *** |
| | | insectDmel | -0.008 | 0.009 | -0.853 | 0.396 | | | insectDmel | 0.64 | 0.194 | 3.3 | **0.002** | ** |
| | | insectSpal | -0.0008 | 0.009 | -0.087 | 0.931 | | | insectSpal | 0.008 | 0.016 | 0.49 | 0.626 | |
| | | conc1 | -0.0002 | <0.001 | -2.209 | **0.03** | * | | conc1 | -0.0004 | <0.001 | -3.314 | **0.002** | ** |
| | | insectDmel:conc1 | 0.00002 | <0.001 | 0.188 | 0.851 | | | insectDmel:conc1 | -0.066 | 0.0194 | -3.414 | **0.001** | ** |
| | | insectSpal:conc1 | 0.00001 | <0.001 | 0.12 | 0.905 | | | insectSpal:conc1 | 0.00006 | <0.001 | 0.326 | 0.745 | |
| | I-a | (Intercept) | 0.106 | 0.016 | 6.599 | **<0.001** | *** | S-b | (Intercept) | 0.037 | 0.005 | 7.034 | **<0.001** | *** |
| | | insectDmel | -0.032 | 0.023 | -1.396 | 0.168 | | | insectDmel | 0.03 | 0.0143 | 2.192 | **0.032** | * |
| | | insectSpal | -0.015 | 0.023 | -0.635 | 0.528 | | | insectSpal | 0.048 | 0.0157 | 3.063 | **0.003** | ** |
| | | conc1 | -0.001 | <0.001 | -3.666 | **0.001** | *** | | conc1 | -0.0002 | <0.001 | -3.323 | **0.002** | ** |
| | | insectDmel:conc1 | 0.0003 | <0.001 | 1.155 | 0.253 | | | insectDmel:conc1 | -0.0003 | <0.001 | -2.021 | **0.048** | * |
| | | insectSpal:conc1 | 0.0003 | <0.001 | 0.89 | 0.377 | | | insectSpal:conc1 | -0.0002 | <0.001 | -1.135 | 0.261 | |
| | I-b | (Intercept) | 0.094 | 0.018 | 5.349 | **<0.001** | *** | S-c | (Intercept) | 0.033 | 0.005 | 5.993 | **<0.001** | *** |
| | | insectDmel | -0.009 | 0.025 | -0.356 | 0.723 | | | insectDmel | 0.012 | 0.011 | 1.108 | 0.273 | |
| | | insectSpal | -0.049 | 0.019 | -2.608 | **0.012** | * | | insectSpal | 0.021 | 0.0111 | 1.861 | 0.069 | . |
| | | conc1 | -0.0007 | <0.001 | -4 | **<0.001** | *** | | conc1 | -0.0002 | <0.001 | -2.246 | **0.029** | * |
| | | insectDmel:conc1 | 0.0001 | <0.001 | 0.366 | 0.716 | | | insectDmel:conc1 | -0.0002 | <0.001 | -1.334 | 0.188 | |
| | | insectSpal:conc1 | 0.0006 | <0.001 | 2.772 | **0.008** | ** | | insectSpal:conc1 | -0.0002 | <0.001 | -1.547 | 0.128 | |

**Table S4.** Summary statistics of electrophysiological responses to glucosinolates (sinigrin, gluconasturtiin, and neoglucobrassicin), using generalized linear models with gamma distributions **(Fig. 2e).** Spikes per second from labellar sensilla was regressed against species, concentration, and their interaction term.

| Compound | Sensilla | | Estimate | Std. Error | t value | Pr(>\|t\|) | | Sensilla | | Estimate | Std. Error | t value | Pr(>\|t\|) | |
|---|---|---|---|---|---|---|---|---|---|---|---|---|---|---|
| sinigrin | L | (Intercept) | 0.897 | 0.159 | 5.628 | **<0.001** | *** | S-a | (Intercept) | 0.117 | 0.023 | 5.092 | **<0.001** | *** |
| | | insectDmel | -0.346 | 0.202 | -1.711 | 0.091 | . | | insectDmel | 0.492 | 0.177 | 2.782 | **0.007** | ** |
| | | insectSpal | 0.123 | 0.255 | 0.48 | 0.632 | | | insectSpal | 0.511 | 0.138 | 3.694 | **<0.001** | *** |
| | | conc1 | -0.026 | 0.005 | -4.729 | **<0.001** | *** | | conc1 | -0.002 | 0.001 | -1.756 | 0.083 | . |
| | | insectDmel:conc1 | 0.022 | 0.009 | 2.464 | **0.016** | * | | insectDmel:conc1 | 0.002 | 0.01 | 0.155 | 0.877 | |
| | | insectSpal:conc1 | 0.017 | 0.013 | 1.319 | 0.19 | | | insectSpal:conc1 | -0.011 | 0.006 | -2.025 | **0.046** | * |
| | I-a | (Intercept) | 0.185 | 0.037 | 4.959 | **<0.001** | *** | S-b | (Intercept) | 0.043 | 0.005 | 8.529 | **<0.001** | *** |
| | | insectDmel | 0.428 | 0.182 | 2.348 | **0.021** | * | | insectDmel | 0.82 | 0.147 | 5.59 | **<0.001** | *** |
| | | insectSpal | -0.018 | 0.051 | -0.358 | 0.721 | | | insectSpal | 0.433 | 0.077 | 5.637 | **<0.001** | *** |
| | | conc1 | -0.002 | 0.002 | -1.391 | 0.168 | | | conc1 | -0.001 | <0.001 | -2.771 | **0.007** | ** |
| | | insectDmel:conc1 | 0.005 | 0.012 | 0.471 | 0.638 | | | insectDmel:conc1 | -0.019 | 0.006 | -3.364 | **0.001** | ** |
| | | insectSpal:conc1 | 0.002 | 0.003 | 0.659 | 0.512 | | | insectSpal:conc1 | -0.006 | 0.003 | -1.738 | 0.086 | . |
| | I-b | (Intercept) | 0.284 | 0.051 | 5.581 | **<0.001** | *** | S-c | (Intercept) | 0.086 | 0.02 | 4.205 | **<0.001** | *** |
| | | insectDmel | 0.515 | 0.187 | 2.762 | **0.007** | ** | | insectDmel | 0.93 | 0.335 | 2.772 | **0.007** | ** |
| | | insectSpal | 0.049 | 0.077 | 0.631 | 0.53 | | | insectSpal | 0.217 | 0.08 | 2.716 | **0.008** | ** |
| | | conc1 | -0.007 | 0.002 | -3.676 | **<0.001** | *** | | conc1 | -0.002 | 0.001 | -2.571 | **0.012** | * |
| | | insectDmel:conc1 | 0.015 | 0.014 | 1.115 | 0.269 | | | insectDmel:conc1 | -0.003 | 0.018 | -0.166 | 0.869 | |
| | | insectSpal:conc1 | 0.002 | 0.003 | 0.454 | 0.651 | | | insectSpal:conc1 | -0.005 | 0.003 | -1.54 | 0.127 | |
| gluconasturtiin | L | (Intercept) | 0.829 | 0.15 | 5.514 | **<0.001** | *** | S-a | (Intercept) | 0.057 | 0.016 | 3.603 | **0.001** | *** |
| | | insectDmel | -0.058 | 0.196 | -0.295 | 0.769 | | | insectDmel | 0.151 | 0.066 | 2.298 | **0.024** | * |
| | | insectSpal | 0.163 | 0.23 | 0.71 | 0.48 | | | insectSpal | 0.142 | 0.052 | 2.732 | **0.008** | ** |
| | | conc1 | 0.002 | 0.01 | 0.232 | 0.817 | | | conc1 | -0.001 | 0.001 | -0.897 | 0.372 | |
| | | insectDmel:conc1 | -0.014 | 0.012 | -1.219 | 0.227 | | | insectDmel:conc1 | 0.008 | 0.007 | 1.127 | 0.263 | |
| | | insectSpal:conc1 | -0.011 | 0.014 | -0.774 | 0.442 | | | insectSpal:conc1 | -0.005 | 0.002 | -2.587 | **0.011** | * |
| | I-a | (Intercept) | 0.153 | 0.028 | 5.549 | **<0.001** | *** | S-b | (Intercept) | 0.051 | 0.011 | 4.733 | **<0.001** | *** |
| | | insectDmel | 0.316 | 0.11 | 2.868 | **0.005** | ** | | insectDmel | 0.228 | 0.071 | 3.222 | **0.002** | ** |
| | | insectSpal | -0.038 | 0.035 | -1.09 | 0.279 | | | insectSpal | 0.223 | 0.056 | 3.983 | **<0.001** | *** |
| | | conc1 | -0.003 | 0.001 | -2.818 | **0.006** | ** | | conc1 | 0.001 | 0.001 | 1.416 | 0.161 | |
| | | insectDmel:conc1 | 0.018 | 0.012 | 1.585 | 0.116 | | | insectDmel:conc1 | 0.014 | 0.009 | 1.576 | 0.119 | |
| | | insectSpal:conc1 | 0.0003 | 0.001 | 0.228 | 0.82 | | | insectSpal:conc1 | -0.004 | 0.003 | -1.277 | 0.205 | |
| | I-b | (Intercept) | 0.104 | 0.015 | 6.757 | **<0.001** | *** | S-c | (Intercept) | 0.054 | 0.015 | 3.497 | **0.001** | *** |

| | | | | | | | | | | | | | | |
|---|---|---|---|---|---|---|---|---|---|---|---|---|---|---|
| | | insectDmel | 0.896 | 0.178 | 5.046 | **<0.001** | *** | | insectDmel | 0.767 | 0.235 | 3.263 | **0.002** | ** |
| | | insectSpal | 0.0004 | 0.023 | 0.016 | 0.987 | | | insectSpal | 0.153 | 0.059 | 2.6 | **0.011** | * |
| | | conc1 | -0.002 | 0.001 | -2.343 | **0.023** | * | | conc1 | 0.004 | 0.002 | 2.009 | **0.048** | * |
| | | insectDmel:conc1 | 0.002 | 0.011 | 0.154 | 0.878 | | | insectDmel:conc1 | -0.011 | 0.012 | -0.934 | 0.353 | |
| | | insectSpal:conc1 | -0.0004 | 0.001 | -0.382 | 0.704 | | | insectSpal:conc1 | 0.013 | 0.009 | 1.426 | 0.158 | |
| neoglucobrassicin | L | (Intercept) | 0.929 | 0.070 | 13.311 | **<0.001** | *** | S-a | (Intercept) | 0.094 | 0.025 | 3.827 | **<0.001** | *** |
| | | insectDmel | 0.071 | 0.100 | 0.718 | 0.476 | | | insectDmel | 0.141 | 0.057 | 2.465 | **0.016** | * |
| | | insectSpal | -0.073 | 0.094 | -0.775 | 0.441 | | | insectSpal | 0.024 | 0.036 | 0.663 | 0.509 | |
| | | conc1 | 0.003 | 0.005 | 0.58 | 0.564 | | | conc1 | -0.002 | 0.001 | -2.133 | **0.036** | * |
| | | insectDmel:conc1 | -0.003 | 0.007 | -0.422 | 0.674 | | | insectDmel:conc1 | -0.003 | 0.002 | -1.494 | 0.139 | |
| | | insectSpal:conc1 | 0.00008 | 0.006 | 0.013 | 0.989 | | | insectSpal:conc1 | -0.001 | 0.001 | -0.728 | 0.469 | |
| | I-a | (Intercept) | 0.159 | 0.036 | 4.417 | **<0.001** | *** | S-b | (Intercept) | 0.075 | 0.024 | 3.073 | **0.003** | ** |
| | | insectDmel | 0.515 | 0.149 | 3.453 | **0.001** | *** | | insectDmel | 0.295 | 0.102 | 2.891 | **0.005** | ** |
| | | insectSpal | -0.08 | 0.039 | -2.037 | **0.046** | * | | insectSpal | 0.07 | 0.054 | 1.306 | 0.195 | |
| | | conc1 | -0.003 | 0.001 | -2.237 | **0.029** | * | | conc1 | 0.001 | 0.001 | 0.407 | 0.685 | |
| | | insectDmel:conc1 | 0.017 | 0.013 | 1.321 | 0.191 | | | insectDmel:conc1 | -0.005 | 0.005 | -1.176 | 0.243 | |
| | | insectSpal:conc1 | 0.001 | 0.001 | 0.749 | 0.456 | | | insectSpal:conc1 | 0.01 | 0.006 | 1.793 | 0.077 | . |
| | I-b | (Intercept) | 0.147 | 0.039 | 3.728 | **<0.001** | *** | S-c | (Intercept) | 0.063 | 0.01 | 6.616 | **<0.001** | *** |
| | | insectDmel | 0.742 | 0.235 | 3.163 | **0.002** | ** | | insectDmel | 0.665 | 0.114 | 5.819 | **<0.001** | *** |
| | | insectSpal | -0.052 | 0.046 | -1.127 | 0.264 | | | insectSpal | 0.02 | 0.019 | 1.052 | 0.296 | |
| | | conc1 | -0.003 | 0.002 | -1.862 | 0.068 | . | | conc1 | -0.001 | <0.001 | -1.039 | 0.302 | |
| | | insectDmel:conc1 | 0.003 | 0.015 | 0.165 | 0.87 | | | insectDmel:conc1 | 0.011 | 0.009 | 1.274 | 0.207 | |
| | | insectSpal:conc1 | 0.001 | 0.002 | 0.395 | 0.694 | | | insectSpal:conc1 | 0.041 | 0.006 | 6.602 | **<0.001** | *** |

**Table S5.** Summary statistics of electrophysiological responses to six glucosinolates. This includes gluconasturtiin, glucotropaeolin, neoglucobrassicin, glucobrassicin, sinigrin, glucoraphanin at 10 mM, using generalized linear models with gamma distributions (**Fig. 2e**). Spikes per second from labellar sensilla was regressed against species, concentration, and their interaction term.

| | | | **Kruskal-Wallis test** | | | **Post-hoc Dunn's test** | | | |
|---|---|---|---|---|---|---|---|---|---|
| **Compound** | **Sensilla** | **Conc. (mM)** | **chi-squared** | **df** | **p-value** | **Comparison** | **Z** | **P.unadj** | **P.adj** |
| gluconasturtiin | I-a | 10 | 11.228 | 2 | **0.004** | Dmel-Sfla | -2.128 | 0.033 | 0.1 |
| | | | | | | Dmel-Spal | -3.346 | 0.001 | **0.002** |
| | | | | | | Sfla-Spal | -1.462 | 0.144 | 0.431 |
| | I-b | 10 | 9.514 | 2 | **0.009** | Dmel-Sfla | -2.534 | 0.011 | **0.034** |
| | | | | | | Dmel-Spal | -2.869 | 0.004 | **0.012** |
| | | | | | | Sfla-Spal | -0.649 | 0.516 | 1 |
| | S-a | 10 | 12.77 | 2 | **0.002** | Dmel-Sfla | -3.411 | 0.001 | **0.002** |
| | | | | | | Dmel-Spal | -1.185 | 0.236 | 0.708 |
| | | | | | | Sfla-Spal | 2.594 | 0.009 | **0.028** |
| glucotropaeolin | I-a | 10 | 14.251 | 2 | **0.0008** | Dmel-Sfla | -1.803 | 0.071 | 0.214 |
| | | | | | | Dmel-Spal | -3.744 | 0 | **0.001** |
| | | | | | | Sfla-Spal | -2.206 | 0.027 | 0.082 |
| | I-b | 10 | 7.9917 | 2 | **0.018** | Dmel-Sfla | -1.078 | 0.281 | 0.843 |
| | | | | | | Dmel-Spal | -2.802 | 0.005 | **0.015** |
| | | | | | | Sfla-Spal | -1.724 | 0.085 | 0.254 |
| | S-a | 10 | 5.606 | 2 | 0.06 | | | | |
| neoglucobrassicin | I-a | 10 | 13.345 | 2 | **0.001** | Dmel-Sfla | -2.109 | 0.035 | 0.105 |
| | | | | | | Dmel-Spal | -3.627 | 0 | **0.001** |
| | | | | | | Sfla-Spal | -1.35 | 0.177 | 0.531 |
| | I-b | 10 | 12.946 | 2 | **0.002** | Dmel-Sfla | -1.923 | 0.054 | 0.163 |
| | | | | | | Dmel-Spal | -3.595 | 0 | **0.001** |
| | | | | | | Sfla-Spal | -1.672 | 0.094 | 0.283 |
| | S-a | 10 | 7.779 | 2 | **0.02** | Dmel-Sfla | -2.391 | 0.017 | 0.05 |
| | | | | | | Dmel-Spal | -2.464 | 0.014 | **0.041** |
| | | | | | | Sfla-Spal | 0.005 | 0.996 | 1 |
| glucobrassicin | I-a | 10 | 9.508 | 2 | **0.009** | Dmel-Sfla | -2.764 | 0.006 | **0.017** |
| | | | | | | Dmel-Spal | -2.695 | 0.007 | **0.021** |
| | | | | | | Sfla-Spal | -0.084 | 0.933 | 1 |
| | I-b | 10 | 11.421 | 2 | **0.003** | Dmel-Sfla | -1.974 | 0.048 | 0.145 |
| | | | | | | Dmel-Spal | -3.369 | 0.001 | **0.002** |
| | | | | | | Sfla-Spal | -1.522 | 0.128 | 0.384 |
| | S-a | 10 | 9.502 | 2 | **0.009** | Dmel-Sfla | -2.917 | 0.004 | **0.011** |
| | | | | | | Dmel-Spal | -2.175 | 0.03 | 0.089 |
| | | | | | | Sfla-Spal | 0.384 | 0.701 | 1 |
| sinigrin | I-a | 10 | 0.35 | 2 | 0.84 | | | | |
| | I-b | 10 | 4.264 | 2 | 0.119 | | | | |
| | S-a | 10 | 9.39 | 2 | **0.009** | Dmel-Sfla | -2.181 | 0.029 | 0.087 |
| | | | | | | Dmel-Spal | -0.651 | 0.515 | 1 |
| | | | | | | Sfla-Spal | 2.625 | 0.009 | **0.026** |
| glucoraphanin | I-a | 10 | 8.245 | 2 | **0.016** | Dmel-Sfla | -2.447 | 0.014 | **0.043** |
| | | | | | | Dmel-Spal | -2.584 | 0.01 | **0.029** |
| | | | | | | Sfla-Spal | -0.143 | 0.886 | 1 |
| | I-b | 10 | 8.818 | 2 | **0.012** | Dmel-Sfla | -2.911 | 0.004 | **0.011** |
| | | | | | | Dmel-Spal | -0.87 | 0.384 | 1 |
| | | | | | | Sfla-Spal | 2.081 | 0.037 | 0.112 |
| | S-a | 10 | 12.518 | 2 | **0.002** | Dmel-Sfla | -3.23 | 0.001 | **0.004** |
| | | | | | | Dmel-Spal | -0.344 | 0.731 | 1 |
| | | | | | | Sfla-Spal | 3.014 | 0.003 | **0.008** |

**Table S6.** Summary statistics of electrophysiological responses towards mixed solutions of sinigrin (1 and 10 mM) and sucrose (5 mM), using Welch two sample t-test **(Fig. S4e)**

| Species | mean (1 mM sinigrin) | mean (10 mM sinigrin) | t | df | p-value | 95% confidence interval |
|---|---|---|---|---|---|---|
| Dmel | 19.733 | 7.571 | 4.148 | 19 | **<0.001** | [6.025, 18.298] |
| Spal | 11.286 | 2.667 | 4.471 | 10 | **0.001** | [4.325, 12.913] |
| Sfla | 18.333 | 11 | 2.107 | 15 | 0.053 | [-0.099, 14.766] |

**Table S7.** Maximum likelihood tests for positive selection from PAML analyses. Abbreviations: “bk” = background, “fr” = foreground, “prop.” = proportion of sites within site class.

| | | | | site class: 0 | | | site class: 1 | | | site class: 2a | | | site class: 2b | | | ln L | 2*ΔlnL | *P* (df = 2) | q-value (5% FDR) |
|---|---|---|---|---|---|---|---|---|---|---|---|---|---|---|---|---|---|---|---|
| Gene | Model | κ (ts/tv) | tree length | prop. | bk ω | fr ω | prop. | bk ω | fr ω | prop. | bk ω | fr ω | prop. | bk ω | fr ω | | | | |
| **Lineage preceding *S. flava* specified as foreground** | | | | | | | | | | | | | | | | | | | |
| *Gr02a* | Branch-site Model A | 1.83 | 5.119 | 0.537 | 0.09034 | 0.09034 | 0.103 | 1 | 1 | 0.302 | 0.09034 | 1 | 0.058 | 1 | 1 | -6328 | -5.80E-05 | 1 | 0.89858793 |
| *Gr02a* | Branch-site Model A (ω = 1) | 1.83 | 5.119 | 0.332 | 0.09033 | 0.09033 | 0.064 | 1 | 1 | 0.507 | 0.09033 | 1 | 0.097 | 1 | 1 | -6328 | | | |
| *Gr05a* | Branch-site Model A | 1.86 | 5.416 | 0.765 | 0.07743 | 0.07743 | 0.235 | 1 | 1 | 0 | 0.07743 | 1 | 0 | 1 | 1 | -7001 | -1344.71 | 1 | 0.89858793 |
| *Gr05a* | Branch-site Model A (ω = 1) | 1.86 | 5.416 | 0.765 | 0.07743 | 0.07743 | 0.235 | 1 | 1 | 0 | 0.07743 | 1 | 0 | 1 | 1 | -7001 | | | |
| *Gr08a* | Branch-site Model A | 1.96 | 6.165 | 0.79 | 0.12346 | 0.12346 | 0.209 | 1 | 1 | 0.001 | 0.12346 | 1 | 0 | 1 | 1 | -6753 | 0 | 1 | 0.89858793 |
| *Gr08a* | Branch-site Model A (ω = 1) | 1.96 | 6.165 | 0.789 | 0.12346 | 0.12346 | 0.209 | 1 | 1 | 0.002 | 0.12346 | 1 | 0 | 1 | 1 | -6753 | | | |
| *Gr09a* | Branch-site Model A | 2.27 | 10.2 | 0.535 | 0.11431 | 0.11431 | 0.165 | 1 | 1 | 0.229 | 0.11431 | 3.55328 | 0.071 | 1 | 3.55 | -6492 | -0.00028 | 1 | 0.89858793 |
| *Gr09a* | Branch-site Model A (ω = 1) | 2.27 | 10.201 | 0.642 | 0.11429 | 0.11429 | 0.198 | 1 | | 0.123 | 0.11429 | 1 | 0.038 | 1 | 1 | -6492 | | | |
| *Gr10a* | Branch-site Model A | 1.44 | 5.983 | 0.676 | 0.07913 | 0.07913 | 0.144 | 1 | 1 | 0.148 | 0.07913 | 1 | 0.032 | 1 | 1 | -6875 | 4.00E-06 | 0.998404 | 0.89858793 |
| *Gr10a* | Branch-site Model A (ω = 1) | 1.44 | 5.983 | 0.642 | 0.07913 | 0.07913 | 0.137 | 1 | 1 | 0.182 | 0.07913 | 1 | 0.039 | 1 | 1 | -6875 | | | |
| *Gr21a* | Branch-site Model A | 1.68 | 3.34 | 0.097 | 0.02459 | 0.02459 | 0.004 | 1 | 1 | 0.867 | 0.02459 | 6.75024 | 0.032 | 1 | 6.75 | -5152 | -0.00305 | 1 | 0.89858793 |
| *Gr21a* | Branch-site Model A (ω = 1) | 1.68 | 3.34 | 0.964 | 0.02459 | 0.02459 | 0.036 | 1 | 1 | 0 | 0.02459 | 1 | 0 | 1 | 1 | -5152 | | | |
| *Gr23a* | Branch-site Model A | 1.9 | 7.655 | 0.713 | 0.14572 | 0.14572 | 0.287 | 1 | 1 | 0 | 0.14572 | 1 | 0 | 1 | 1 | -12802 | 0 | 1 | 0.89858793 |
| *Gr23a* | Branch-site Model A (ω = 1) | 1.9 | 7.655 | 0.697 | 0.14572 | 0.14572 | 0.28 | 1 | 1 | 0.017 | 0.14572 | 1 | 0.007 | 1 | 1 | -12802 | | | |
| *Gr28a* | Branch-site Model A | 1.75 | 3.586 | 0 | 0.03357 | 0.03357 | 0 | 1 | 1 | 0.936 | 0.03357 | 1 | 0.064 | 1 | 1 | -5544 | 0 | 1 | 0.89858793 |
| *Gr28a* | Branch-site Model A (ω = 1) | 1.75 | 3.586 | 0 | 0.03357 | 0.03357 | 0 | 1 | 1 | 0.936 | 0.03357 | 1 | 0.064 | 1 | 1 | -5544 | | | |
| *Gr28bB* | Branch-site Model A | 1.94 | 3.057 | 0.001 | 0.03015 | 0.03015 | 0 | 1 | 1 | 0.949 | 0.03015 | 2.26587 | 0.05 | 1 | 2.27 | -5307 | 6.20E-05 | 0.993718 | 0.89858793 |
| *Gr28bB* | Branch-site Model A (ω = 1) | 1.94 | 3.057 | 0 | 0.03015 | 0.03015 | 0 | 1 | 1 | 0.95 | 0.03015 | 1 | 0.05 | 1 | 1 | -5307 | | | |
| *Gr28bC* | Branch-site Model A | 1.74 | 5.724 | 0.836 | 0.06101 | 0.06101 | 0.153 | 1 | 1 | 0.01 | 0.06101 | 1 | 0.002 | 1 | 1 | -9578 | -8542.64 | 1 | 0.89858793 |
| *Gr28bC* | Branch-site Model A (ω = 1) | 1.74 | 5.724 | 0.836 | 0.06101 | 0.06101 | 0.153 | 1 | 1 | 0.01 | 0.06101 | 1 | 0.002 | 1 | 1 | -9578 | | | |

| | | | | | | | | | | | | | | | | | | | |
|---|---|---|---|---|---|---|---|---|---|---|---|---|---|---|---|---|---|---|---|
| *Gr28bD* | Branch-site Model<br>A | 1.82 | 4.006 | 0.00<br>1 | 0.0507<br>6 | 0.0507<br>6 | 0 | 1 | 1 | 0.89<br>3 | 0.0507<br>6 | 68.4172<br>8 | 0.10<br>6 | 1 | 68.<br>4 | -6068 | 0.01793<br>6 | 0.89346<br>2 | 0.8985879<br>3 |
| *Gr28bD* | Branch-site Model<br>A (ω = 1) | 1.82 | 4.006 | 0.37<br>5 | 0.0507<br>9 | 0.0507<br>9 | 0.04<br>5 | 1 | 1 | 0.51<br>8 | 0.0507<br>9 | 1 | 0.06<br>2 | 1 | 1 | -6068 | | | |
| *Gr28bE* | Branch-site Model<br>A | 2.11 | 4.891 | 0.70<br>6 | 0.0633<br>6 | 0.0633<br>6 | 0.25 | 1 | 1 | 0.03<br>2 | 0.0633<br>6 | 999 | 0.01<br>1 | 1 | 999 | -7094 | 4.59196<br>6 | 0.03212<br>2 | 0.2040236 |
| *Gr28bE* | Branch-site Model<br>A (ω = 1) | 2.09 | 4.881 | 0.63<br>7 | 0.0633<br>1 | 0.0633<br>1 | 0.22<br>6 | 1 | 1 | 0.10<br>1 | 0.0633<br>1 | 1 | 0.03<br>6 | 1 | 1 | -7096 | | | |
| *Gr32a* | Branch-site Model<br>A | 1.73 | 4.226 | 0.73 | 0.0694 | 0.0694 | 0.10<br>9 | 1 | 1 | 0.14 | 0.0694 | 2.38968 | 0.02<br>1 | 1 | 2.3<br>9 | -6582 | 8.00E-<br>06 | 0.99774<br>3 | 0.8985879<br>3 |
| *Gr32a* | Branch-site Model<br>A (ω = 1) | 1.73 | 4.226 | 0.82<br>3 | 0.0694 | 0.0694 | 0.12<br>3 | 1 | 1 | 0.04<br>7 | 0.0694 | 1 | 0.00<br>7 | 1 | 1 | -6582 | | | |
| *Gr33a* | Branch-site Model<br>A | 1.81 | 3.227 | 0.00<br>4 | 0.0520<br>7 | 0.0520<br>7 | 0.00<br>1 | 1 | 1 | 0.84<br>5 | 0.0520<br>7 | 1 | 0.15<br>1 | 1 | 1 | -6064 | -5.20E-<br>05 | 1 | 0.8985879<br>3 |
| *Gr33a* | Branch-site Model<br>A (ω = 1) | 1.81 | 3.227 | 0 | 0.0520<br>7 | 0.0520<br>7 | 0 | 1 | 1 | 0.84<br>9 | 0.0520<br>7 | 1 | 0.15<br>1 | 1 | 1 | -6064 | | | |
| *Gr39aB* | Branch-site Model<br>A | 1.96 | 6.152 | 0.54<br>9 | 0.0806<br>5 | 0.0806<br>5 | 0.14<br>6 | 1 | 1 | 0.24<br>1 | 0.0806<br>5 | 1 | 0.06<br>4 | 1 | 1 | -6176 | -223.319 | 1 | 0.8985879<br>3 |
| *Gr39aB* | Branch-site Model<br>A (ω = 1) | 1.96 | 6.152 | 0.54<br>9 | 0.0806<br>5 | 0.0806<br>5 | 0.14<br>6 | 1 | 1 | 0.24<br>1 | 0.0806<br>5 | 1 | 0.06<br>4 | 1 | 1 | -6176 | | | |
| *Gr39aD* | Branch-site Model<br>A | 1.95 | 6.096 | 0.50<br>5 | 0.0807<br>8 | 0.0807<br>8 | 0.28<br>6 | 1 | 1 | 0.13<br>3 | 0.0807<br>8 | 1 | 0.07<br>6 | 1 | 1 | -6434 | -0.01669 | 1 | 0.8985879<br>3 |
| *Gr39aD* | Branch-site Model<br>A (ω = 1) | 1.95 | 6.096 | 0.61<br>9 | 0.0807<br>5 | 0.0807<br>5 | 0.35<br>1 | 1 | 1 | 0.01<br>9 | 0.0807<br>5 | 1 | 0.01<br>1 | 1 | 1 | -6434 | | | |
| *Gr39b* | Branch-site Model<br>A | 2.19 | 6.72 | 0.84<br>6 | 0.1011<br>9 | 0.1011<br>9 | 0.15<br>4 | 1 | 1 | 0 | 0.1011<br>9 | 1 | 0 | 1 | 1 | -5142 | 2583.97<br>7 | 1 | 0.8985879<br>3 |
| *Gr39b* | Branch-site Model<br>A (ω = 1) | 2.19 | 6.72 | 0.84<br>6 | 0.1011<br>9 | 0.1011<br>9 | 0.15<br>4 | 1 | 1 | 0 | 0.1011<br>9 | 1 | 0 | 1 | 1 | -5142 | | | |
| *Gr43a* | Branch-site Model<br>A | 1.66 | 4.613 | 0.31<br>3 | 0.0502<br>8 | 0.0502<br>8 | 0.01<br>7 | 1 | 1 | 0.63<br>7 | 0.0502<br>8 | 4.82692 | 0.03<br>4 | 1 | 4.8<br>3 | -5832 | -0.00263 | 1 | 0.8985879<br>3 |
| *Gr43a* | Branch-site Model<br>A (ω = 1) | 1.66 | 4.613 | 0.94<br>9 | 0.0502<br>7 | 0.0502<br>7 | 0.05 | 1 | 1 | 0.00<br>1 | 0.0502<br>7 | 1 | 0 | 1 | 1 | -5832 | | | |
| *Gr47b* | Branch-site Model<br>A | 1.73 | 6.797 | 0.70<br>8 | 0.1245<br>8 | 0.1245<br>8 | 0.27<br>2 | 1 | 1 | 0.01<br>5 | 0.1245<br>8 | 999 | 0.00<br>6 | 1 | 999 | -8212 | 12.825 | 0.00034<br>2 | 0.0063003<br>3 |
| *Gr47b* | Branch-site Model<br>A (ω = 1) | 1.72 | 6.774 | 0.55<br>9 | 0.1253<br>5 | 0.1253<br>5 | 0.21<br>2 | 1 | 1 | 0.16<br>7 | 0.1253<br>5 | 1 | 0.06<br>3 | 1 | 1 | -8219 | | | |
| *Gr57a* | Branch-site Model<br>A | 2.08 | 6.535 | 0.83<br>6 | 0.1246<br>5 | 0.1246<br>5 | 0.16<br>4 | 1 | 1 | 0 | 0.1246<br>5 | 1 | 0 | 1 | 1 | -7290 | 0 | 1 | 0.8985879<br>3 |
| *Gr57a* | Branch-site Model<br>A (ω = 1) | 2.08 | 6.535 | 0.83<br>6 | 0.1246<br>5 | 0.1246<br>5 | 0.16<br>4 | 1 | 1 | 0 | 0.1246<br>5 | 1 | 0 | 1 | 1 | -7290 | | | |
| *Gr58b* | Branch-site Model<br>A | 2.13 | 11.41<br>1 | 0.71<br>3 | 0.1487<br>2 | 0.1487<br>2 | 0.25<br>7 | 1 | 1 | 0.02<br>2 | 0.1487<br>2 | 999 | 0.00<br>8 | 1 | 999 | -7734 | 56.1386<br>8 | 6.75E-<br>14 | 2.49E-12 |
| *Gr58b* | Branch-site Model<br>A (ω = 1) | 2.09 | 8.462 | 0.51<br>9 | 0.1474<br>1 | 0.1474<br>1 | 0.17<br>9 | 1 | 1 | 0.22<br>5 | 0.1474<br>1 | 1 | 0.07<br>7 | 1 | 1 | -7762 | | | |
| *Gr58c* | Branch-site Model<br>A | 1.89 | 10.45<br>8 | 0.79 | 0.1065<br>2 | 0.1065<br>2 | 0.15<br>2 | 1 | 1 | 0.04<br>9 | 0.1065<br>2 | 1.41205 | 0.00<br>9 | 1 | 1.4<br>1 | -7038 | 1448.31<br>3 | 0.86989 | 0.8985879<br>3 |
| *Gr58c* | Branch-site Model<br>A (ω = 1) | 1.89 | 10.45<br>7 | 0.77<br>3 | 0.1065<br>3 | 0.1065<br>3 | 0.14<br>8 | 1 | 1 | 0.06<br>6 | 0.1065<br>3 | 1 | 0.01<br>3 | 1 | 1 | -7038 | | | |
| *Gr59a* | Branch-site Model<br>A | 1.71 | 16.04<br>1 | 0.65<br>1 | 0.1912<br>3 | 0.1912<br>3 | 0.28<br>4 | 1 | 1 | 0.04<br>5 | 0.1912<br>3 | 1 | 0.02 | 1 | 1 | -<br>1587<br>4 | -17671.6 | 1 | 0.8985879<br>3 |
| *Gr59a* | Branch-site Model<br>A (ω = 1) | 1.71 | 16.04<br>1 | 0.65<br>1 | 0.1912<br>3 | 0.1912<br>3 | 0.28<br>4 | 1 | 1 | 0.04<br>5 | 0.1912<br>3 | 1 | 0.02 | 1 | 1 | -<br>1587<br>4 | | | |
| *Gr59d* | Branch-site Model<br>A | 2.02 | 28.03<br>7 | 0.66 | 0.2313<br>3 | 0.2313<br>3 | 0.30<br>1 | 1 | 1 | 0.02<br>7 | 0.2313<br>3 | 3.29691 | 0.01<br>2 | 1 | 3.3 | -<br>3010<br>6 | -28464.2 | 0.17909<br>7 | 0.6598316<br>1 |

| | | | | | | | | | | | | | | | | | | | |
|---|---|---|---|---|---|---|---|---|---|---|---|---|---|---|---|---|---|---|---|
| *Gr59d* | Branch-site Model A (ω = 1) | 2.02 | 28.036 | 0.618 | 0.23121 | 0.23121 | 0.281 | 1 | 1 | 0.069 | 0.23121 | 1 | 0.031 | 1 | 1 | -30107 | | | |
| *Gr61a* | Branch-site Model A | 2.22 | 6.936 | 0.843 | 0.07238 | 0.07238 | 0.149 | 1 | 1 | 0.006 | 0.07238 | 999 | 0.001 | 1 | 999 | -6768 | 7.64702 | 0.005687 | 0.06983566 |
| *Gr61a* | Branch-site Model A (ω = 1) | 2.21 | 6.946 | 0.766 | 0.07179 | 0.07179 | 0.136 | 1 | 1 | 0.084 | 0.07179 | 1 | 0.015 | 1 | 1 | -6772 | | | |
| *Gr63a* | Branch-site Model A | 1.76 | 3.549 | 0.272 | 0.03343 | 0.03343 | 0.025 | 1 | 1 | 0.644 | 0.03343 | 5.18374 | 0.059 | 1 | 5.18 | -6114 | -0.00049 | 1 | 0.89858793 |
| *Gr63a* | Branch-site Model A (ω = 1) | 1.76 | 3.549 | 0.916 | 0.03343 | 0.03343 | 0.084 | 1 | 1 | 0 | 0.03343 | 1 | 0 | 1 | 1 | -6114 | | | |
| *Gr64a* | Branch-site Model A | 2.18 | 4.94 | 0 | 0.0776 | 0.0776 | 0 | 1 | 1 | 0.792 | 0.0776 | 1 | 0.208 | 1 | 1 | -7110 | 0 | 1 | 0.89858793 |
| *Gr64a* | Branch-site Model A (ω = 1) | 2.18 | 4.94 | 0 | 0.0776 | 0.0776 | 0 | 1 | 1 | 0.792 | 0.0776 | 1 | 0.208 | 1 | 1 | -7110 | | | |
| *Gr64b* | Branch-site Model A | 1.89 | 3.982 | 0.91 | 0.06083 | 0.06083 | 0.09 | 1 | 1 | 0 | 0.06083 | 1 | 0 | 1 | 1 | -5477 | 0 | 1 | 0.89858793 |
| *Gr64b* | Branch-site Model A (ω = 1) | 1.89 | 3.982 | 0.91 | 0.06083 | 0.06083 | 0.09 | 1 | 1 | 0 | 0.06083 | 1 | 0 | 1 | 1 | -5477 | | | |
| *Gr64c* | Branch-site Model A | 2.06 | 4.595 | 0.903 | 0.08102 | 0.08102 | 0.087 | 1 | 1 | 0.008 | 0.08102 | 999 | 0.001 | 1 | 999 | -6064 | 2.890212 | 0.089119 | 0.41041736 |
| *Gr64c* | Branch-site Model A (ω = 1) | 2.05 | 4.586 | 0.847 | 0.08077 | 0.08077 | 0.083 | 1 | 1 | 0.063 | 0.08077 | 1 | 0.006 | 1 | 1 | -6065 | | | |
| *Gr64e* | Branch-site Model A | 2.07 | 4.948 | 0.849 | 0.062 | 0.062 | 0.151 | 1 | 1 | 0 | 0.062 | 1 | 0 | 1 | 1 | -6671 | 8.00E-06 | 0.997743 | 0.89858793 |
| *Gr64e* | Branch-site Model A (ω = 1) | 2.07 | 4.948 | 0.846 | 0.062 | 0.062 | 0.15 | 1 | 1 | 0.003 | 0.062 | 1 | 0.001 | 1 | 1 | -6671 | | | |
| *Gr64f* | Branch-site Model A | 1.88 | 4.613 | 0 | 0.07473 | 0.07473 | 0 | 1 | 1 | 0.856 | 0.07473 | 999 | 0.144 | 1 | 999 | -6876 | 0.617404 | 0.432014 | 0.89858793 |
| *Gr64f* | Branch-site Model A (ω = 1) | 1.87 | 4.612 | 0 | 0.07472 | 0.07472 | 0 | 1 | 1 | 0.856 | 0.07472 | 1 | 0.144 | 1 | 1 | -6876 | | | |
| *Gr66a* | Branch-site Model A | 1.77 | 4.444 | 0.886 | 0.04636 | 0.04636 | 0.111 | 1 | 1 | 0.003 | 0.04636 | 25.60598 | 0 | 1 | 25.6 | -7373 | 3.472718 | 0.062389 | 0.32836168 |
| *Gr66a* | Branch-site Model A (ω = 1) | 1.76 | 4.447 | 0.84 | 0.04639 | 0.04639 | 0.103 | 1 | 1 | 0.051 | 0.04639 | 1 | 0.006 | 1 | 1 | -7374 | | | |
| *Gr77a* | Branch-site Model A | 1.49 | 7.596 | 0.652 | 0.16063 | 0.16063 | 0.269 | 1 | 1 | 0.056 | 0.16063 | 1 | 0.023 | 1 | 1 | -8224 | -1698.58 | 1 | 0.89858793 |
| *Gr77a* | Branch-site Model A (ω = 1) | 1.49 | 7.596 | 0.652 | 0.16063 | 0.16063 | 0.269 | 1 | 1 | 0.056 | 0.16063 | 1 | 0.023 | 1 | 1 | -8224 | | | |
| *Gr85a* | Branch-site Model A | 1.89 | 27.37 | 0.548 | 0.26939 | 0.26939 | 0.452 | 1 | 1 | 0 | 0.26939 | 1 | 0 | 1 | 1 | -26367 | 0 | 1 | 0.89858793 |
| *Gr85a* | Branch-site Model A (ω = 1) | 1.89 | 27.37 | 0.548 | 0.26939 | 0.26939 | 0.452 | 1 | 1 | 0 | 0.26939 | 1 | 0 | 1 | 1 | -26367 | | | |
| *Gr89a* | Branch-site Model A | 2.15 | 5.542 | 0 | 0.08555 | 0.08555 | 0 | 1 | 1 | 0.845 | 0.08555 | 1 | 0.155 | 1 | 1 | -5022 | 0 | 1 | 0.89858793 |
| *Gr89a* | Branch-site Model A (ω = 1) | 2.15 | 5.542 | 0 | 0.08555 | 0.08555 | 0 | 1 | 1 | 0.845 | 0.08555 | 1 | 0.155 | 1 | 1 | -5022 | | | |
| *Gr93a* | Branch-site Model A | 1.72 | 6.825 | 0.795 | 0.09588 | 0.09588 | 0.183 | 1 | 1 | 0.018 | 0.09588 | 999 | 0.004 | 1 | 999 | -7216 | 0.291978 | 0.588956 | 0.89858793 |
| *Gr93a* | Branch-site Model A (ω = 1) | 1.72 | 6.831 | 0.725 | 0.09561 | 0.09561 | 0.167 | 1 | 1 | 0.088 | 0.09561 | 1 | 0.02 | 1 | 1 | -7216 | | | |
| *Gr93c* | Branch-site Model A | 1.93 | 12.264 | 0.633 | 0.1711 | 0.1711 | 0.227 | 1 | 1 | 0.103 | 0.1711 | 2.42675 | 0.037 | 1 | 2.43 | -14124 | -13815.7 | 0.137955 | 0.56472906 |

| | | | | | | | | | | | | | | | | | | | |
|---|---|---|---|---|---|---|---|---|---|---|---|---|---|---|---|---|---|---|---|
| *Gr93c* | Branch-site Model A (ω = 1) | 1.93 | 12.256 | 0.57 | 0.17125 | 0.17125 | 0.205 | 1 | 1 | 0.166 | 0.17125 | 1 | 0.06 | 1 | 1 | -14125 | | | |
| *Gr94a* | Branch-site Model A | 1.82 | 7.835 | 0 | 0.11583 | 0.11583 | 0 | 1 | 1 | 0.887 | 0.11583 | 1 | 0.113 | 1 | 1 | -7503 | 0 | 1 | 0.89858793 |
| *Gr94a* | Branch-site Model A (ω = 1) | 1.82 | 7.835 | 0 | 0.11583 | 0.11583 | 0 | 1 | 1 | 0.887 | 0.11583 | 1 | 0.113 | 1 | 1 | -7503 | | | |
| *Gr97a* | Branch-site Model A | 1.98 | 6.832 | 0 | 0.14857 | 0.14857 | 0 | 1 | 1 | 0.79 | 0.14857 | 1 | 0.21 | 1 | 1 | -7823 | 0 | 1 | 0.89858793 |
| *Gr97a* | Branch-site Model A (ω = 1) | 1.98 | 6.832 | 0 | 0.14857 | 0.14857 | 0 | 1 | 1 | 0.79 | 0.14857 | 1 | 0.21 | 1 | 1 | -7823 | | | |
| *Gr98a* | Branch-site Model A | 2 | 16.928 | 0.729 | 0.21901 | 0.21901 | 0.268 | 1 | 1 | 0.002 | 0.21901 | 22.14829 | 0.001 | 1 | 22.1 | -19596 | -23545.5 | 0.027665 | 0.2040236 |
| *Gr98a* | Branch-site Model A (ω = 1) | 2 | 16.95 | 0.626 | 0.21659 | 0.21659 | 0.229 | 1 | 1 | 0.107 | 0.21659 | 1 | 0.039 | 1 | 1 | -19598 | | | |
| *Gr98bcd* | Branch-site Model A | 2.09 | 23.51 | 0.718 | 0.16279 | 0.16279 | 0.273 | 1 | 1 | 0.006 | 0.16279 | 15.73784 | 0.002 | 1 | 15.7 | -12241 | 14714.11 | 0.033227 | 0.2040236 |
| *Gr98bcd* | Branch-site Model A (ω = 1) | 2.08 | 23.477 | 0.721 | 0.16291 | 0.16291 | 0.273 | 1 | 1 | 0.004 | 0.16291 | 1 | 0.002 | 1 | 1 | -12243 | | | |
| **Lineage preceding mustard feeders (*S. flava* and *S. montana*) specified as foreground** | | | | | | | | | | | | | | | | | | | |
| *Gr02a* | Branch-site Model A | 1.83 | 5.117 | 0.839 | 0.09044 | 0.09044 | 0.161 | 1 | 1 | 0 | 0.09044 | 1 | 0 | 1 | 1 | -6328 | 0 | 1 | 0.947696 |
| *Gr02a* | Branch-site Model A (ω = 1) | 1.83 | 5.117 | 0.839 | 0.09044 | 0.09044 | 0.161 | 1 | 1 | 0 | 0.09044 | 1 | 0 | 1 | 1 | -6328 | | | |
| *Gr05a* | Branch-site Model A | 1.86 | 5.51 | 0.768 | 0.07859 | 0.07859 | 0.217 | 1 | 1 | 0.012 | 0.07859 | 75.6173 | 0.003 | 1 | 75.6 | -6997 | 6.32 | 0.011938 | 0.181583 |
| *Gr05a* | Branch-site Model A (ω = 1) | 1.86 | 5.441 | 0.706 | 0.0774 | 0.0774 | 0.211 | 1 | 1 | 0.063 | 0.0774 | 1 | 0.019 | 1 | 1 | -7000 | | | |
| *Gr08a* | Branch-site Model A | 1.96 | 6.165 | 0.791 | 0.12346 | 0.12346 | 0.209 | 1 | 1 | 0 | 0.12346 | 1 | 0 | 1 | 1 | -6753 | 0 | 1 | 0.947696 |
| *Gr08a* | Branch-site Model A (ω = 1) | 1.96 | 6.165 | 0.791 | 0.12346 | 0.12346 | 0.209 | 1 | 1 | 0 | 0.12346 | 1 | 0 | 1 | 1 | -6753 | | | |
| *Gr09a* | Branch-site Model A | 2.27 | 10.2 | 0.764 | 0.11432 | 0.11432 | 0.236 | 1 | 1 | 0 | 0.11432 | 1 | 0 | 1 | 1 | -6492 | 0 | 1 | 0.947696 |
| *Gr09a* | Branch-site Model A (ω = 1) | 2.27 | 10.2 | 0.764 | 0.11432 | 0.11432 | 0.236 | 1 | 1 | 0 | 0.11432 | 1 | 0 | 1 | 1 | -6492 | | | |
| *Gr10a* | Branch-site Model A | 1.45 | 6.046 | 0.753 | 0.07651 | 0.07651 | 0.162 | 1 | 1 | 0.07 | 0.07651 | 1.27441 | 0.015 | 1 | 1.27 | -6874 | 0.04 | 0.841481 | 0.947696 |
| *Gr10a* | Branch-site Model A (ω = 1) | 1.45 | 6.045 | 0.738 | 0.0765 | 0.0765 | 0.159 | 1 | 1 | 0.085 | 0.0765 | 1 | 0.018 | 1 | 1 | -6874 | | | |
| *Gr21a* | Branch-site Model A | 1.68 | 3.359 | 0.91 | 0.02397 | 0.02397 | 0.033 | 1 | 1 | 0.055 | 0.02397 | 1 | 0.002 | 1 | 1 | -5150 | 0 | 1 | 0.947696 |
| *Gr21a* | Branch-site Model A (ω = 1) | 1.68 | 3.359 | 0.91 | 0.02397 | 0.02397 | 0.033 | 1 | 1 | 0.055 | 0.02397 | 1 | 0.002 | 1 | 1 | -5150 | | | |
| *Gr23a* | Branch-site Model A | 1.91 | 7.742 | 0.584 | 0.14097 | 0.14097 | 0.233 | 1 | 1 | 0.131 | 0.14097 | 2.15141 | 0.052 | 1 | 2.15 | -12798 | 1 | 0.317311 | 0.947696 |
| *Gr23a* | Branch-site Model A (ω = 1) | 1.9 | 7.738 | 0.495 | 0.14089 | 0.14089 | 0.198 | 1 | 1 | 0.219 | 0.14089 | 1 | 0.088 | 1 | 1 | -12798 | | | |
| *Gr28a* | Branch-site Model A | 1.76 | 3.626 | 0 | 0.03246 | 0.03246 | 0 | 1 | 1 | 0.936 | 0.03246 | 1 | 0.064 | 1 | 1 | -5551 | -15.8 | 1 | 0.947696 |

| | | | | | | | | | | | | | | | | | | | |
|---|---|---|---|---|---|---|---|---|---|---|---|---|---|---|---|---|---|---|---|
| *Gr28a* | Branch-site Model A (ω = 1) | 1.75 | 3.598 | 0.866 | 0.03307 | 0.03307 | 0.059 | 1 | 1 | 0.07 | 0.03307 | 1 | 0.005 | 1 | 1 | -5543 | | | |
| *Gr28bB* | Branch-site Model A | 1.94 | 3.075 | 0.726 | 0.02882 | 0.02882 | 0.038 | 1 | 1 | 0.225 | 0.02882 | 1.46939 | 0.012 | 1 | 1.47 | -5302 | 0.02 | 0.887537 | 0.947696 |
| *Gr28bB* | Branch-site Model A (ω = 1) | 1.94 | 3.075 | 0.636 | 0.02881 | 0.02881 | 0.033 | 1 | 1 | 0.314 | 0.02881 | 1 | 0.016 | 1 | 1 | -5302 | | | |
| *Gr28bC* | Branch-site Model A | 1.74 | 5.762 | 0.588 | 0.05633 | 0.05633 | 0.107 | 1 | 1 | 0.258 | 0.05633 | 1 | 0.047 | 1 | 1 | -9564 | 0 | 1 | 0.947696 |
| *Gr28bC* | Branch-site Model A (ω = 1) | 1.74 | 5.762 | 0.588 | 0.05633 | 0.05633 | 0.107 | 1 | 1 | 0.258 | 0.05633 | 1 | 0.047 | 1 | 1 | -9564 | | | |
| *Gr28bD* | Branch-site Model A | 1.82 | 4.035 | 0.884 | 0.04953 | 0.04953 | 0.103 | 1 | 1 | 0.012 | 0.04953 | 11.67236 | 0.001 | 1 | 11.7 | -6063 | 2.52 | 0.112411 | 0.683932 |
| *Gr28bD* | Branch-site Model A (ω = 1) | 1.82 | 4.031 | 0.8 | 0.04945 | 0.04945 | 0.093 | 1 | 1 | 0.096 | 0.04945 | 1 | 0.011 | 1 | 1 | -6065 | | | |
| *Gr28bE* | Branch-site Model A | 2.1 | 4.849 | 0.728 | 0.06412 | 0.06412 | 0.265 | 1 | 1 | 0.005 | 0.06412 | 1 | 0.002 | 1 | 1 | -7099 | 0 | 1 | 0.947696 |
| *Gr28bE* | Branch-site Model A (ω = 1) | 2.1 | 4.849 | 0.728 | 0.06412 | 0.06412 | 0.265 | 1 | 1 | 0.005 | 0.06412 | 1 | 0.002 | 1 | 1 | -7099 | | | |
| *Gr32a* | Branch-site Model A | 1.74 | 4.275 | 0.698 | 0.06491 | 0.06491 | 0.105 | 1 | 1 | 0.171 | 0.06491 | 1 | 0.026 | 1 | 1 | -6577 | 0 | 1 | 0.947696 |
| *Gr32a* | Branch-site Model A (ω = 1) | 1.74 | 4.275 | 0.698 | 0.06491 | 0.06491 | 0.105 | 1 | 1 | 0.171 | 0.06491 | 1 | 0.026 | 1 | 1 | -6577 | | | |
| *Gr33a* | Branch-site Model A | 1.82 | 3.247 | 0.72 | 0.04911 | 0.04911 | 0.128 | 1 | 1 | 0.13 | 0.04911 | 1.26659 | 0.023 | 1 | 1.27 | -6060 | 0.04 | 0.841481 | 0.947696 |
| *Gr33a* | Branch-site Model A (ω = 1) | 1.82 | 3.246 | 0.693 | 0.04907 | 0.04907 | 0.123 | 1 | 1 | 0.157 | 0.04907 | 1 | 0.028 | 1 | 1 | -6060 | | | |
| *Gr39aA* | Branch-site Model A | 2.01 | 27.803 | 0.503 | 0.12683 | 0.12683 | 0.493 | 1 | 1 | 0.002 | 0.12683 | 13.70702 | 0.002 | 1 | 13.7 | -31859 | 30.8 | 2.86E-08 | 8.70E-06 |
| *Gr39aA* | Branch-site Model A (ω = 1) | 2 | 27.781 | 0.508 | 0.1291 | 0.1291 | 0.492 | 1 | 1 | 0 | 0.1291 | 1 | 0 | 1 | 1 | -31874 | | | |
| *Gr39aB* | Branch-site Model A | 1.96 | 5.888 | 0.676 | 0.09514 | 0.09514 | 0.182 | 1 | 1 | 0.112 | 0.09514 | 4.59989 | 0.03 | 1 | 4.6 | -6182 | 4.66 | 0.030873 | 0.347848 |
| *Gr39aB* | Branch-site Model A (ω = 1) | 1.94 | 5.866 | 0.467 | 0.09439 | 0.09439 | 0.127 | 1 | 1 | 0.318 | 0.09439 | 1 | 0.087 | 1 | 1 | -6184 | | | |
| *Gr39aD* | Branch-site Model A | 1.96 | 6.147 | 0.635 | 0.08151 | 0.08151 | 0.347 | 1 | 1 | 0.012 | 0.08151 | 21.22866 | 0.006 | 1 | 21.2 | -6431 | 4.36 | 0.036792 | 0.361049 |
| *Gr39aD* | Branch-site Model A (ω = 1) | 1.95 | 6.139 | 0.529 | 0.08123 | 0.08123 | 0.29 | 1 | 1 | 0.118 | 0.08123 | 1 | 0.064 | 1 | 1 | -6433 | | | |
| *Gr39b* | Branch-site Model A | 2.19 | 6.77 | 0.827 | 0.10059 | 0.10059 | 0.14 | 1 | 1 | 0.028 | 0.10059 | 7.89758 | 0.005 | 1 | 7.9 | -5141 | 1.34 | 0.247034 | 0.947696 |
| *Gr39b* | Branch-site Model A (ω = 1) | 2.18 | 6.768 | 0.766 | 0.10004 | 0.10004 | 0.134 | 1 | 1 | 0.085 | 0.10004 | 1 | 0.015 | 1 | 1 | -5142 | | | |
| *Gr43a* | Branch-site Model A | 1.66 | 4.639 | 0.899 | 0.04938 | 0.04938 | 0.047 | 1 | 1 | 0.052 | 0.04938 | 1 | 0.003 | 1 | 1 | -5831 | 0 | 1 | 0.947696 |
| *Gr43a* | Branch-site Model A (ω = 1) | 1.66 | 4.639 | 0.899 | 0.04938 | 0.04938 | 0.047 | 1 | 1 | 0.052 | 0.04938 | 1 | 0.003 | 1 | 1 | -5831 | | | |
| *Gr47b* | Branch-site Model A | 1.74 | 6.834 | 0.466 | 0.1189 | 0.1189 | 0.185 | 1 | 1 | 0.249 | 0.1189 | 1 | 0.099 | 1 | 1 | -8216 | 0 | 1 | 0.947696 |
| *Gr47b* | Branch-site Model A (ω = 1) | 1.74 | 6.834 | 0.466 | 0.1189 | 0.1189 | 0.185 | 1 | 1 | 0.249 | 0.1189 | 1 | 0.099 | 1 | 1 | -8216 | | | |
| *Gr57a* | Branch-site Model A | 2.09 | 6.6 | 0.82 | 0.12172 | 0.12172 | 0.161 | 1 | 1 | 0.016 | 0.12172 | 13.17626 | 0.003 | 1 | 13.2 | -7286 | 3.52 | 0.060632 | 0.533403 |
| *Gr57a* | Branch-site Model A (ω = 1) | 2.08 | 6.589 | 0.718 | 0.12163 | 0.12163 | 0.14 | 1 | 1 | 0.119 | 0.12163 | 1 | 0.023 | 1 | 1 | -7288 | | | |
| *Gr58b* | Branch-site Model A | 2.08 | 8.279 | 0.69 | 0.15408 | 0.15408 | 0.249 | 1 | 1 | 0.045 | 0.15408 | 1.81947 | 0.016 | 1 | 1.82 | -7770 | 0.08 | 0.777297 | 0.947696 |

| | | | | | | | | | | | | | | | | | | | |
|---|---|---|---|---|---|---|---|---|---|---|---|---|---|---|---|---|---|---|---|
| *Gr58b* | Branch-site Model A (ω = 1) | 2.08 | 8.276 | 0.662 | 0.15412 | 0.15412 | 0.239 | 1 | 1 | 0.072 | 0.15412 | 1 | 0.026 | 1 | 1 | -7770 | | | |
| *Gr58c* | Branch-site Model A | 1.89 | 10.452 | 0.832 | 0.10675 | 0.10675 | 0.162 | 1 | 1 | 0.005 | 0.10675 | 16.97855 | 0.001 | 1 | 17 | -7038 | 1.78 | 0.182149 | 0.801826 |
| *Gr58c* | Branch-site Model A (ω = 1) | 1.89 | 10.402 | 0.831 | 0.10747 | 0.10747 | 0.161 | 1 | 1 | 0.007 | 0.10747 | 1 | 0.001 | 1 | 1 | -7039 | | | |
| *Gr59a* | Branch-site Model A | 1.71 | 16.101 | 0.611 | 0.189 | 0.189 | 0.263 | 1 | 1 | 0.088 | 0.189 | 1.83028 | 0.038 | 1 | 1.83 | -15871 | 0.4 | 0.527089 | 0.947696 |
| *Gr59a* | Branch-site Model A (ω = 1) | 1.71 | 16.101 | 0.549 | 0.1887 | 0.1887 | 0.237 | 1 | 1 | 0.15 | 0.1887 | 1 | 0.065 | 1 | 1 | -15872 | | | |
| *Gr59d* | Branch-site Model A | 2.01 | 28.036 | 0.661 | 0.2319 | 0.2319 | 0.297 | 1 | 1 | 0.029 | 0.2319 | 1.34396 | 0.013 | 1 | 1.34 | -30108 | 0 | 1 | 0.947696 |
| *Gr59d* | Branch-site Model A (ω = 1) | 2.01 | 28.036 | 0.647 | 0.23174 | 0.23174 | 0.291 | 1 | 1 | 0.042 | 0.23174 | 1 | 0.019 | 1 | 1 | -30108 | | | |
| *Gr61a* | Branch-site Model A | 2.22 | 6.868 | 0.846 | 0.0734 | 0.0734 | 0.154 | 1 | 1 | 0 | 0.0734 | 1 | 0 | 1 | 1 | -6774 | 0 | 1 | 0.947696 |
| *Gr61a* | Branch-site Model A (ω = 1) | 2.22 | 6.868 | 0.846 | 0.0734 | 0.0734 | 0.154 | 1 | 1 | 0 | 0.0734 | 1 | 0 | 1 | 1 | -6774 | | | |
| *Gr63a* | Branch-site Model A | 1.76 | 3.549 | 0.916 | 0.03343 | 0.03343 | 0.084 | 1 | 1 | 0 | 0.03343 | 1 | 0 | 1 | 1 | -6114 | 0 | 1 | 0.947696 |
| *Gr63a* | Branch-site Model A (ω = 1) | 1.76 | 3.549 | 0.916 | 0.03343 | 0.03343 | 0.084 | 1 | 1 | 0 | 0.03343 | 1 | 0 | 1 | 1 | -6114 | | | |
| *Gr64a* | Branch-site Model A | 2.18 | 4.923 | 0.791 | 0.07835 | 0.07835 | 0.209 | 1 | 1 | 0 | 0.07835 | 1 | 0 | 1 | 1 | -7110 | 0 | 1 | 0.947696 |
| *Gr64a* | Branch-site Model A (ω = 1) | 2.18 | 4.923 | 0.791 | 0.07835 | 0.07835 | 0.209 | 1 | 1 | 0 | 0.07835 | 1 | 0 | 1 | 1 | -7110 | | | |
| *Gr64b* | Branch-site Model A | 1.88 | 4.004 | 0.897 | 0.06001 | 0.06001 | 0.083 | 1 | 1 | 0.018 | 0.06001 | 5.91284 | 0.002 | 1 | 5.91 | -5475 | 1.46 | 0.22693 | 0.932899 |
| *Gr64b* | Branch-site Model A (ω = 1) | 1.88 | 4.001 | 0.839 | 0.0597 | 0.0597 | 0.079 | 1 | 1 | 0.075 | 0.0597 | 1 | 0.007 | 1 | 1 | -5475 | | | |
| *Gr64c* | Branch-site Model A | 2.07 | 4.602 | 0.782 | 0.07905 | 0.07905 | 0.079 | 1 | 1 | 0.126 | 0.07905 | 1 | 0.013 | 1 | 1 | -6064 | 0 | 1 | 0.947696 |
| *Gr64c* | Branch-site Model A (ω = 1) | 2.07 | 4.602 | 0.782 | 0.07905 | 0.07905 | 0.079 | 1 | 1 | 0.126 | 0.07905 | 1 | 0.013 | 1 | 1 | -6064 | | | |
| *Gr64e* | Branch-site Model A | 2.07 | 4.948 | 0.849 | 0.062 | 0.062 | 0.151 | 1 | 1 | 0 | 0.062 | 1 | 0 | 1 | 1 | -6671 | 0 | 1 | 0.947696 |
| *Gr64e* | Branch-site Model A (ω = 1) | 2.07 | 4.948 | 0.849 | 0.062 | 0.062 | 0.151 | 1 | 1 | 0 | 0.062 | 1 | 0 | 1 | 1 | -6671 | | | |
| *Gr64f* | Branch-site Model A | 1.87 | 4.615 | 0.844 | 0.07596 | 0.07596 | 0.139 | 1 | 1 | 0.015 | 0.07596 | 10.80759 | 0.002 | 1 | 10.8 | -6876 | 2.04 | 0.15321 | 0.745971 |
| *Gr64f* | Branch-site Model A (ω = 1) | 1.87 | 4.608 | 0.769 | 0.07571 | 0.07571 | 0.128 | 1 | 1 | 0.088 | 0.07571 | 1 | 0.015 | 1 | 1 | -6877 | | | |
| *Gr66a* | Branch-site Model A | 1.76 | 4.474 | 0.852 | 0.04557 | 0.04557 | 0.103 | 1 | 1 | 0.04 | 0.04557 | 5.20025 | 0.005 | 1 | 5.2 | -7372 | 1.02 | 0.312519 | 0.947696 |
| *Gr66a* | Branch-site Model A (ω = 1) | 1.76 | 4.472 | 0.727 | 0.04563 | 0.04563 | 0.087 | 1 | 1 | 0.166 | 0.04563 | 1 | 0.02 | 1 | 1 | -7372 | | | |
| *Gr77a* | Branch-site Model A | 1.5 | 7.618 | 0.669 | 0.16009 | 0.16009 | 0.271 | 1 | 1 | 0.043 | 0.16009 | 3.63659 | 0.017 | 1 | 3.64 | -8223 | 0.58 | 0.446312 | 0.947696 |
| *Gr77a* | Branch-site Model A (ω = 1) | 1.49 | 7.607 | 0.629 | 0.16017 | 0.16017 | 0.257 | 1 | 1 | 0.081 | 0.16017 | 1 | 0.033 | 1 | 1 | -8223 | | | |
| *Gr85a* | Branch-site Model A | 1.89 | 27.381 | 0.501 | 0.26901 | 0.26901 | 0.412 | 1 | 1 | 0.048 | 0.26901 | 7.69249 | 0.039 | 1 | 7.69 | -26367 | 0.8 | 0.371093 | 0.947696 |

| | | | | | | | | | | | | | | | | | | |
|---|---|---|---|---|---|---|---|---|---|---|---|---|---|---|---|---|---|---|
| *Gr85a* | Branch-site Model A (ω = 1) | 1.89 | 27.37 | 0.548 | 0.26939 | 0.26939 | 0.452 | 1 | 1 | 0 | 0.26939 | 1 | 0 | 1 | 1 | -26367 | | |
| *Gr89a* | Branch-site Model A | 2.15 | 5.559 | 0.841 | 0.08685 | 0.08685 | 0.146 | 1 | 1 | 0.011 | 0.08685 | 12.88904 | 0.002 | 1 | 12.9 | -5022 | 1.4 | 0.236724 | 0.947696 |
| *Gr89a* | Branch-site Model A (ω = 1) | 2.15 | 5.52 | 0.827 | 0.08674 | 0.08674 | 0.149 | 1 | 1 | 0.021 | 0.08674 | 1 | 0.004 | 1 | 1 | -5023 | | |
| *Gr93a* | Branch-site Model A | 1.72 | 6.837 | 0.791 | 0.09569 | 0.09569 | 0.176 | 1 | 1 | 0.027 | 0.09569 | 9.49844 | 0.006 | 1 | 9.5 | -7214 | 2.34 | 0.12609 | 0.730337 |
| *Gr93a* | Branch-site Model A (ω = 1) | 1.72 | 6.83 | 0.673 | 0.09576 | 0.09576 | 0.15 | 1 | 1 | 0.145 | 0.09576 | 1 | 0.032 | 1 | 1 | -7215 | | |
| *Gr93c* | Branch-site Model A | 1.93 | 12.277 | 0.599 | 0.17141 | 0.17141 | 0.208 | 1 | 1 | 0.143 | 0.17141 | 2.03121 | 0.05 | 1 | 2.03 | -14124 | 2 | 0.157299 | 0.745971 |
| *Gr93c* | Branch-site Model A (ω = 1) | 1.92 | 12.268 | 0.548 | 0.1715 | 0.1715 | 0.191 | 1 | 1 | 0.193 | 0.1715 | 1 | 0.067 | 1 | 1 | -14125 | | |
| *Gr94a* | Branch-site Model A | 1.83 | 8.014 | 0.532 | 0.1094 | 0.1094 | 0.067 | 1 | 1 | 0.356 | 0.1094 | 1 | 0.045 | 1 | 1 | -7496 | 0 | 1 | 0.947696 |
| *Gr94a* | Branch-site Model A (ω = 1) | 1.83 | 8.014 | 0.532 | 0.1094 | 0.1094 | 0.067 | 1 | 1 | 0.356 | 0.1094 | 1 | 0.045 | 1 | 1 | -7496 | | |
| *Gr97a* | Branch-site Model A | 1.98 | 6.837 | 0.733 | 0.14704 | 0.14704 | 0.2 | 1 | 1 | 0.053 | 0.14704 | 1 | 0.014 | 1 | 1 | -7823 | 0 | 1 | 0.947696 |
| *Gr97a* | Branch-site Model A (ω = 1) | 1.98 | 6.837 | 0.733 | 0.14704 | 0.14704 | 0.2 | 1 | 1 | 0.053 | 0.14704 | 1 | 0.014 | 1 | 1 | -7823 | | |
| *Gr98a* | Branch-site Model A | 1.99 | 17.019 | 0.689 | 0.2173 | 0.2173 | 0.229 | 1 | 1 | 0.062 | 0.2173 | 4.82749 | 0.02 | 1 | 4.83 | -19590 | 11.2 | 0.000818 | 0.035549 |
| *Gr98a* | Branch-site Model A (ω = 1) | 1.99 | 16.993 | 0.598 | 0.21639 | 0.21639 | 0.206 | 1 | 1 | 0.146 | 0.21639 | 1 | 0.05 | 1 | 1 | -19596 | | |
| *Gr98bcd* | Branch-site Model A | 2.08 | 23.631 | 0.707 | 0.16109 | 0.16109 | 0.263 | 1 | 1 | 0.021 | 0.16109 | 17.50494 | 0.008 | 1 | 17.5 | -12239 | 2.6 | 0.106864 | 0.683932 |
| *Gr98bcd* | Branch-site Model A (ω = 1) | 2.08 | 23.654 | 0.411 | 0.16077 | 0.16077 | 0.153 | 1 | 1 | 0.318 | 0.16077 | 1 | 0.118 | 1 | 1 | -12241 | | |

**Lineage preceding all herbivores specified as foreground (from Pelaez et al. 2023)**

| | | | | | | | | | | | | | | | | | | |
|---|---|---|---|---|---|---|---|---|---|---|---|---|---|---|---|---|---|---|
| *Gr02a* | Branch-site Model A | 1.84 | 5.207 | 0.833 | 0.08958 | 0.08958 | 0.159 | 1 | 1 | 0.008 | 0.08958 | 56.45923 | 0.001 | 1 | 56.5 | -6326 | 5.1 | 0.023926 | 0.290156 |
| *Gr02a* | Branch-site Model A (ω = 1) | 1.83 | 5.126 | 0.828 | 0.09006 | 0.09006 | 0.158 | 1 | 1 | 0.011 | 0.09006 | 1 | 0.002 | 1 | 1 | -6328 | | |
| *Gr05a* | Branch-site Model A | 1.86 | 5.416 | 0.765 | 0.07743 | 0.07743 | 0.235 | 1 | 1 | 0 | 0.07743 | 1 | 0 | 1 | 1 | -7001 | 0 | 1 | 0.949005 |
| *Gr05a* | Branch-site Model A (ω = 1) | 1.86 | 5.416 | 0.765 | 0.07743 | 0.07743 | 0.235 | 1 | 1 | 0 | 0.07743 | 1 | 0 | 1 | 1 | -7001 | | |
| *Gr08a* | Branch-site Model A | 1.96 | 6.165 | 0.791 | 0.12346 | 0.12346 | 0.209 | 1 | 1 | 0 | 0.12346 | 1 | 0 | 1 | 1 | -6753 | 0 | 1 | 0.949005 |
| *Gr08a* | Branch-site Model A (ω = 1) | 1.96 | 6.165 | 0.791 | 0.12346 | 0.12346 | 0.209 | 1 | 1 | 0 | 0.12346 | 1 | 0 | 1 | 1 | -6753 | | |
| *Gr09a* | Branch-site Model A | 2.27 | 10.2 | 0.764 | 0.11432 | 0.11432 | 0.236 | 1 | 1 | 0 | 0.11432 | 1 | 0 | 1 | 1 | -6492 | 0 | 1 | 0.949005 |
| *Gr09a* | Branch-site Model A (ω = 1) | 2.27 | 10.2 | 0.764 | 0.11432 | 0.11432 | 0.236 | 1 | 1 | 0 | 0.11432 | 1 | 0 | 1 | 1 | -6492 | | |
| *Gr10a* | Branch-site Model A | 1.44 | 5.998 | 0.824 | 0.07891 | 0.07891 | 0.172 | 1 | 1 | 0.004 | 0.07891 | 10.5712 | 0.001 | 1 | 10.6 | -6875 | 0.82 | 0.36518 | 0.949005 |

| | | | | | | | | | | | | | | | | | | | |
|---|---|---|---|---|---|---|---|---|---|---|---|---|---|---|---|---|---|---|---|
| *Gr10a* | Branch-site Model A (ω = 1) | 1.44 | 5.983 | 0.824 | 0.07913 | 0.07913 | 0.176 | 1 | 1 | 0 | 0.07913 | 1 | 0 | 1 | 1 | -6875 | | | |
| *Gr21a* | Branch-site Model A | 1.68 | 3.35 | 0.957 | 0.02402 | 0.02402 | 0.036 | 1 | 1 | 0.006 | 0.02402 | 3.07374 | 0 | 1 | 3.07 | -5151 | 0.16 | 0.689157 | 0.949005 |
| *Gr21a* | Branch-site Model A (ω = 1) | 1.68 | 3.349 | 0.949 | 0.02397 | 0.02397 | 0.036 | 1 | 1 | 0.015 | 0.02397 | 1 | 0.001 | 1 | 1 | -5151 | | | |
| *Gr23a* | Branch-site Model A | 1.9 | 7.693 | 0.708 | 0.14403 | 0.14403 | 0.284 | 1 | 1 | 0.006 | 0.14403 | 8.21015 | 0.002 | 1 | 8.21 | -12801 | 2.4 | 0.121335 | 0.683799 |
| *Gr23a* | Branch-site Model A (ω = 1) | 1.9 | 7.685 | 0.685 | 0.14398 | 0.14398 | 0.274 | 1 | 1 | 0.029 | 0.14398 | 1 | 0.011 | 1 | 1 | -12802 | | | |
| *Gr28a* | Branch-site Model A | 1.75 | 3.575 | 0.936 | 0.0343 | 0.0343 | 0.064 | 1 | 1 | 0 | 0.0343 | 1 | 0 | 1 | 1 | -5545 | 0 | 1 | 0.949005 |
| *Gr28a* | Branch-site Model A (ω = 1) | 1.75 | 3.575 | 0.936 | 0.0343 | 0.0343 | 0.064 | 1 | 1 | 0 | 0.0343 | 1 | 0 | 1 | 1 | -5545 | | | |
| *Gr28bA* | Branch-site Model A | 1.95 | 7.756 | 0 | 0.04638 | 0.04638 | 0 | 1 | 1 | 0.667 | 0.04638 | 1 | 0.333 | 1 | 1 | -7335 | 0 | 1 | 0.949005 |
| *Gr28bA* | Branch-site Model A (ω = 1) | 1.95 | 7.756 | 0 | 0.04638 | 0.04638 | 0 | 1 | 1 | 0.667 | 0.04638 | 1 | 0.333 | 1 | 1 | -7335 | | | |
| *Gr28bB* | Branch-site Model A | 1.94 | 3.055 | 0.951 | 0.03046 | 0.03046 | 0.049 | 1 | 1 | 0 | 0.03046 | 1 | 0 | 1 | 1 | -5307 | 0 | 1 | 0.949005 |
| *Gr28bB* | Branch-site Model A (ω = 1) | 1.94 | 3.055 | 0.951 | 0.03046 | 0.03046 | 0.049 | 1 | 1 | 0 | 0.03046 | 1 | 0 | 1 | 1 | -5307 | | | |
| *Gr28bC* | Branch-site Model A | 1.74 | 5.724 | 0.845 | 0.06116 | 0.06116 | 0.153 | 1 | 1 | 0.002 | 0.06116 | 3.88689 | 0 | 1 | 3.89 | -9578 | 0.16 | 0.689157 | 0.949005 |
| *Gr28bC* | Branch-site Model A (ω = 1) | 1.74 | 5.723 | 0.845 | 0.06115 | 0.06115 | 0.154 | 1 | 1 | 0.001 | 0.06115 | 1 | 0 | 1 | 1 | -9578 | | | |
| *Gr28bD* | Branch-site Model A | 1.82 | 4.005 | 0.894 | 0.05092 | 0.05092 | 0.106 | 1 | 1 | 0 | 0.05092 | 1 | 0 | 1 | 1 | -6068 | 0 | 1 | 0.949005 |
| *Gr28bD* | Branch-site Model A (ω = 1) | 1.82 | 4.005 | 0.894 | 0.05092 | 0.05092 | 0.106 | 1 | 1 | 0 | 0.05092 | 1 | 0 | 1 | 1 | -6068 | | | |
| *Gr28bE* | Branch-site Model A | 2.1 | 4.848 | 0.733 | 0.06426 | 0.06426 | 0.267 | 1 | 1 | 0 | 0.06426 | 1 | 0 | 1 | 1 | -7099 | 0 | 1 | 0.949005 |
| *Gr28bE* | Branch-site Model A (ω = 1) | 2.1 | 4.848 | 0.733 | 0.06426 | 0.06426 | 0.267 | 1 | 1 | 0 | 0.06426 | 1 | 0 | 1 | 1 | -7099 | | | |
| *Gr32a* | Branch-site Model A | 1.73 | 4.226 | 0.87 | 0.0694 | 0.0694 | 0.13 | 1 | 1 | 0 | 0.0694 | 1 | 0 | 1 | 1 | -6582 | 0 | 1 | 0.949005 |
| *Gr32a* | Branch-site Model A (ω = 1) | 1.73 | 4.226 | 0.87 | 0.0694 | 0.0694 | 0.13 | 1 | 1 | 0 | 0.0694 | 1 | 0 | 1 | 1 | -6582 | | | |
| *Gr33a* | Branch-site Model A | 1.82 | 3.246 | 0.76 | 0.04808 | 0.04808 | 0.138 | 1 | 1 | 0.087 | 0.04808 | 1 | 0.016 | 1 | 1 | -6061 | 0 | 1 | 0.949005 |
| *Gr33a* | Branch-site Model A (ω = 1) | 1.82 | 3.246 | 0.76 | 0.04808 | 0.04808 | 0.138 | 1 | 1 | 0.087 | 0.04808 | 1 | 0.016 | 1 | 1 | -6061 | | | |
| *Gr39aA* | Branch-site Model A | 2 | 27.787 | 0.494 | 0.12812 | 0.12812 | 0.48 | 1 | 1 | 0.014 | 0.12812 | 1 | 0.013 | 1 | 1 | -31873 | 1 | 0.317311 | 0.949005 |
| *Gr39aA* | Branch-site Model A (ω = 1) | 2 | 27.781 | 0.508 | 0.1291 | 0.1291 | 0.492 | 1 | 1 | 0 | 0.1291 | 1 | 0 | 1 | 1 | -31874 | | | |
| *Gr39aB* | Branch-site Model A | 1.94 | 5.678 | 0.774 | 0.1047 | 0.1047 | 0.226 | 1 | 1 | 0 | 0.1047 | 1 | 0 | 1 | 1 | -6196 | 0 | 1 | 0.949005 |
| *Gr39aB* | Branch-site Model A (ω = 1) | 1.94 | 5.678 | 0.774 | 0.1047 | 0.1047 | 0.226 | 1 | 1 | 0 | 0.1047 | 1 | 0 | 1 | 1 | -6196 | | | |
| *Gr39aD* | Branch-site Model A | 1.95 | 6.093 | 0.638 | 0.08124 | 0.08124 | 0.362 | 1 | 1 | 0 | 0.08124 | 1 | 0 | 1 | 1 | -6434 | 0 | 1 | 0.949005 |
| *Gr39aD* | Branch-site Model A (ω = 1) | 1.95 | 6.093 | 0.638 | 0.08124 | 0.08124 | 0.362 | 1 | 1 | 0 | 0.08124 | 1 | 0 | 1 | 1 | -6434 | | | |

| | | | | | | | | | | | | | | | | | | | |
|---|---|---|---|---|---|---|---|---|---|---|---|---|---|---|---|---|---|---|---|
| *Gr39b* | Branch-site Model A | 2.19 | 6.72 | 0.84 6 | 0.1011 9 | 0.1011 9 | 0.15 4 | 1 | 1 | 0 | 0.1011 9 | 1 | 0 | 1 | 1 | -5142 | 0 | 1 | 0.949005 |
| *Gr39b* | Branch-site Model A (ω = 1) | 2.19 | 6.72 | 0.84 6 | 0.1011 9 | 0.1011 9 | 0.15 4 | 1 | 1 | 0 | 0.1011 9 | 1 | 0 | 1 | 1 | -5142 | | | |
| *Gr43a* | Branch-site Model A | 1.66 | 4.638 | 0.94 7 | 0.0494 4 | 0.0494 4 | 0.05 | 1 | 1 | 0.00 3 | 0.0494 4 | 11.1385 5 | 0 | 1 | 11. 1 | -5830 | 2.66 | 0.10290 1 | 0.652433 |
| *Gr43a* | Branch-site Model A (ω = 1) | 1.66 | 4.634 | 0.93 8 | 0.0494 9 | 0.0494 9 | 0.05 | 1 | 1 | 0.01 2 | 0.0494 9 | 1 | 0.00 1 | 1 | 1 | -5831 | | | |
| *Gr47b* | Branch-site Model A | 1.72 | 6.796 | 0.72 | 0.1270 4 | 0.1270 4 | 0.27 1 | 1 | 1 | 0.00 6 | 0.1270 4 | 21.8720 6 | 0.00 2 | 1 | 21. 9 | -8220 | 2.94 | 0.08641 1 | 0.59724 |
| *Gr47b* | Branch-site Model A (ω = 1) | 1.72 | 6.751 | 0.70 9 | 0.1275 2 | 0.1275 2 | 0.27 | 1 | 1 | 0.01 5 | 0.1275 2 | 1 | 0.00 6 | 1 | 1 | -8221 | | | |
| *Gr57a* | Branch-site Model A | 2.08 | 6.579 | 0.78 3 | 0.1222 5 | 0.1222 5 | 0.15 3 | 1 | 1 | 0.05 3 | 0.1222 5 | 1 | 0.01 | 1 | 1 | -7290 | 0 | 1 | 0.949005 |
| *Gr57a* | Branch-site Model A (ω = 1) | 2.08 | 6.579 | 0.78 3 | 0.1222 5 | 0.1222 5 | 0.15 3 | 1 | 1 | 0.05 3 | 0.1222 5 | 1 | 0.01 | 1 | 1 | -7290 | | | |
| *Gr58b* | Branch-site Model A | 2.1 | 8.358 | 0.72 8 | 0.1549 | 0.1549 | 0.25 6 | 1 | 1 | 0.01 2 | 0.1549 | 20.8037 2 | 0.00 4 | 1 | 20. 8 | -7768 | 3.66 | 0.05573 4 | 0.497957 |
| *Gr58b* | Branch-site Model A (ω = 1) | 2.08 | 8.253 | 0.73 3 | 0.1552 | 0.1552 | 0.26 7 | 1 | 1 | 0 | 0.1552 | 1 | 0 | 1 | 1 | -7770 | | | |
| *Gr58c* | Branch-site Model A | 1.89 | 10.38 9 | 0.83 8 | 0.1076 8 | 0.1076 8 | 0.16 2 | 1 | 1 | 0 | 0.1076 8 | 1 | 0 | 1 | 1 | -7039 | 0 | 1 | 0.949005 |
| *Gr58c* | Branch-site Model A (ω = 1) | 1.89 | 10.38 9 | 0.83 8 | 0.1076 8 | 0.1076 8 | 0.16 2 | 1 | 1 | 0 | 0.1076 8 | 1 | 0 | 1 | 1 | -7039 | | | |
| *Gr59a* | Branch-site Model A | 1.71 | 16.07 1 | 0.69 2 | 0.1921 2 | 0.1921 2 | 0.29 2 | 1 | 1 | 0.01 1 | 0.1921 2 | 5.55213 | 0.00 5 | 1 | 5.5 5 | - 1587 3 | 1.8 | 0.17971 2 | 0.866202 |
| *Gr59a* | Branch-site Model A (ω = 1) | 1.71 | 16.06 4 | 0.66 7 | 0.1912 2 | 0.1912 2 | 0.28 4 | 1 | 1 | 0.03 4 | 0.1912 2 | 1 | 0.01 4 | 1 | 1 | - 1587 4 | | | |
| *Gr59d* | Branch-site Model A | 2.01 | 28.03 6 | 0.68 3 | 0.2323 7 | 0.2323 7 | 0.30 6 | 1 | 1 | 0.00 8 | 0.2323 7 | 3.11604 | 0.00 3 | 1 | 3.1 2 | - 3010 8 | 1 | 0.31731 1 | 0.949005 |
| *Gr59d* | Branch-site Model A (ω = 1) | 2.01 | 28.02 5 | 0.67 7 | 0.2323 4 | 0.2323 4 | 0.30 6 | 1 | 1 | 0.01 2 | 0.2323 4 | 1 | 0.00 5 | 1 | 1 | - 3010 8 | | | |
| *Gr59e* | Branch-site Model A | 2.22 | 8.813 | 0.61 2 | 0.1217 3 | 0.1217 3 | 0.22 5 | 1 | 1 | 0.11 9 | 0.1217 3 | 999 | 0.04 4 | 1 | 999 | -6583 | 4.32 | 0.03766 7 | 0.383278 |
| *Gr59e* | Branch-site Model A (ω = 1) | 2.2 | 8.812 | 0.45 1 | 0.1196 3 | 0.1196 3 | 0.17 | 1 | 1 | 0.27 6 | 0.1196 3 | 1 | 0.10 4 | 1 | 1 | -6586 | | | |
| *Gr59f* | Branch-site Model A | 2.09 | 8.584 | 0.68 3 | 0.1307 7 | 0.1307 7 | 0.27 3 | 1 | 1 | 0.03 2 | 0.1307 7 | 999 | 0.01 3 | 1 | 999 | -7155 | 0.02 | 0.88753 7 | 0.949005 |
| *Gr59f* | Branch-site Model A (ω = 1) | 2.09 | 8.599 | 0.64 9 | 0.1299 6 | 0.1299 6 | 0.26 1 | 1 | 1 | 0.06 4 | 0.1299 6 | 1 | 0.02 6 | 1 | 1 | -7155 | | | |
| *Gr61a* | Branch-site Model A | 2.22 | 6.868 | 0.84 6 | 0.0734 | 0.0734 | 0.15 4 | 1 | 1 | 0 | 0.0734 | 1 | 0 | 1 | 1 | -6774 | 0 | 1 | 0.949005 |
| *Gr61a* | Branch-site Model A (ω = 1) | 2.22 | 6.868 | 0.84 6 | 0.0734 | 0.0734 | 0.15 4 | 1 | 1 | 0 | 0.0734 | 1 | 0 | 1 | 1 | -6774 | | | |
| *Gr63a* | Branch-site Model A | 1.76 | 3.584 | 0.80 8 | 0.0313 9 | 0.0313 9 | 0.07 3 | 1 | 1 | 0.10 9 | 0.0313 9 | 1 | 0.01 | 1 | 1 | -6110 | 16.52 | 4.81E- 05 | 0.009561 |
| *Gr63a* | Branch-site Model A (ω = 1) | 1.77 | 3.605 | 0 | 0.0308 4 | 0.0308 4 | 0 | 1 | 1 | 0.91 8 | 0.0308 4 | 1 | 0.08 2 | 1 | 1 | -6119 | | | |
| *Gr64a* | Branch-site Model A | 2.18 | 4.923 | 0.79 1 | 0.0783 5 | 0.0783 5 | 0.20 9 | 1 | 1 | 0 | 0.0783 5 | 1 | 0 | 1 | 1 | -7110 | 0 | 1 | 0.949005 |
| *Gr64a* | Branch-site Model A (ω = 1) | 2.18 | 4.923 | 0.79 1 | 0.0783 5 | 0.0783 5 | 0.20 9 | 1 | 1 | 0 | 0.0783 5 | 1 | 0 | 1 | 1 | -7110 | | | |
| *Gr64b* | Branch-site Model A | 1.89 | 3.982 | 0.91 | 0.0608 3 | 0.0608 3 | 0.09 | 1 | 1 | 0 | 0.0608 3 | 1 | 0 | 1 | 1 | -5477 | 0 | 1 | 0.949005 |

| | | | | | | | | | | | | | | | | | | | |
|---|---|---|---|---|---|---|---|---|---|---|---|---|---|---|---|---|---|---|---|
| *Gr64b* | Branch-site Model A (ω = 1) | 1.89 | 3.982 | 0.91 | 0.06083 | 0.06083 | 0.09 | 1 | 1 | 0 | 0.06083 | 1 | 0 | 1 | 1 | -5477 | | | |
| *Gr64c* | Branch-site Model A | 2.06 | 4.565 | 0.906 | 0.0812 | 0.0812 | 0.094 | 1 | 1 | 0 | 0.0812 | 1 | 0 | 1 | 1 | -6066 | 0 | 1 | 0.949005 |
| *Gr64c* | Branch-site Model A (ω = 1) | 2.06 | 4.565 | 0.906 | 0.0812 | 0.0812 | 0.094 | 1 | 1 | 0 | 0.0812 | 1 | 0 | 1 | 1 | -6066 | | | |
| *Gr64e* | Branch-site Model A | 2.07 | 4.948 | 0.849 | 0.062 | 0.062 | 0.151 | 1 | 1 | 0 | 0.062 | 1 | 0 | 1 | 1 | -6671 | 0 | 1 | 0.949005 |
| *Gr64e* | Branch-site Model A (ω = 1) | 2.07 | 4.948 | 0.849 | 0.062 | 0.062 | 0.151 | 1 | 1 | 0 | 0.062 | 1 | 0 | 1 | 1 | -6671 | | | |
| *Gr64f* | Branch-site Model A | 1.87 | 4.611 | 0.852 | 0.07585 | 0.07585 | 0.145 | 1 | 1 | 0.002 | 0.07585 | 27.3581 | 0 | 1 | 27.4 | -6877 | 2.64 | 0.104204 | 0.652433 |
| *Gr64f* | Branch-site Model A (ω = 1) | 1.87 | 4.599 | 0.839 | 0.07576 | 0.07576 | 0.144 | 1 | 1 | 0.015 | 0.07576 | 1 | 0.003 | 1 | 1 | -6878 | | | |
| *Gr66a* | Branch-site Model A | 1.77 | 4.459 | 0.825 | 0.04503 | 0.04503 | 0.103 | 1 | 1 | 0.064 | 0.04503 | 1 | 0.008 | 1 | 1 | -7375 | 0 | 1 | 0.949005 |
| *Gr66a* | Branch-site Model A (ω = 1) | 1.77 | 4.459 | 0.825 | 0.04503 | 0.04503 | 0.103 | 1 | 1 | 0.064 | 0.04503 | 1 | 0.008 | 1 | 1 | -7375 | | | |
| *Gr77a* | Branch-site Model A | 1.49 | 7.578 | 0.533 | 0.16185 | 0.16185 | 0.22 | 1 | 1 | 0.175 | 0.16185 | 2.95439 | 0.072 | 1 | 2.95 | -8224 | 0 | 1 | 0.949005 |
| *Gr77a* | Branch-site Model A (ω = 1) | 1.49 | 7.578 | 0.588 | 0.16185 | 0.16185 | 0.242 | 1 | 1 | 0.12 | 0.16185 | 1 | 0.05 | 1 | 1 | -8224 | | | |
| *Gr85a* | Branch-site Model A | 1.89 | 27.38 | 0.543 | 0.2692 | 0.2692 | 0.447 | 1 | 1 | 0.006 | 0.2692 | 5.21371 | 0.005 | 1 | 5.21 | -26367 | 0 | 1 | 0.949005 |
| *Gr85a* | Branch-site Model A (ω = 1) | 1.89 | 27.371 | 0.544 | 0.26932 | 0.26932 | 0.448 | 1 | 1 | 0.004 | 0.26932 | 1 | 0.004 | 1 | 1 | -26367 | | | |
| *Gr89a* | Branch-site Model A | 2.15 | 5.601 | 0.832 | 0.08466 | 0.08466 | 0.146 | 1 | 1 | 0.019 | 0.08466 | 6.26457 | 0.003 | 1 | 6.26 | -5021 | 0.82 | 0.36518 | 0.949005 |
| *Gr89a* | Branch-site Model A (ω = 1) | 2.15 | 5.607 | 0.748 | 0.0842 | 0.0842 | 0.132 | 1 | 1 | 0.102 | 0.0842 | 1 | 0.018 | 1 | 1 | -5022 | | | |
| *Gr93a* | Branch-site Model A | 1.72 | 6.768 | 0.812 | 0.09767 | 0.09767 | 0.188 | 1 | 1 | 0 | 0.09767 | 1 | 0 | 1 | 1 | -7217 | 0 | 1 | 0.949005 |
| *Gr93a* | Branch-site Model A (ω = 1) | 1.72 | 6.768 | 0.812 | 0.09767 | 0.09767 | 0.188 | 1 | 1 | 0 | 0.09767 | 1 | 0 | 1 | 1 | -7217 | | | |
| *Gr93c* | Branch-site Model A | 1.92 | 12.209 | 0.737 | 0.17533 | 0.17533 | 0.263 | 1 | 1 | 0 | 0.17533 | 1 | 0 | 1 | 1 | -14129 | 0 | 1 | 0.949005 |
| *Gr93c* | Branch-site Model A (ω = 1) | 1.92 | 12.209 | 0.737 | 0.17533 | 0.17533 | 0.263 | 1 | 1 | 0 | 0.17533 | 1 | 0 | 1 | 1 | -14129 | | | |
| *Gr94a* | Branch-site Model A | 1.82 | 7.854 | 0.814 | 0.11461 | 0.11461 | 0.105 | 1 | 1 | 0.072 | 0.11461 | 1 | 0.009 | 1 | 1 | -7503 | 0 | 1 | 0.949005 |
| *Gr94a* | Branch-site Model A (ω = 1) | 1.82 | 7.854 | 0.814 | 0.11461 | 0.11461 | 0.105 | 1 | 1 | 0.072 | 0.11461 | 1 | 0.009 | 1 | 1 | -7503 | | | |
| *Gr97a* | Branch-site Model A | 1.98 | 6.814 | 0.785 | 0.14872 | 0.14872 | 0.215 | 1 | 1 | 0 | 0.14872 | 655.2293 | 0 | 1 | 655 | -7823 | -0.68 | 1 | 0.949005 |
| *Gr97a* | Branch-site Model A (ω = 1) | 1.98 | 6.844 | 0.721 | 0.14686 | 0.14686 | 0.193 | 1 | 1 | 0.068 | 0.14686 | 1 | 0.018 | 1 | 1 | -7823 | | | |
| *Gr98a* | Branch-site Model A | 2 | 16.922 | 0.732 | 0.2189 | 0.2189 | 0.268 | 1 | 1 | 0 | 0.2189 | 1 | 0 | 1 | 1 | -19599 | 0 | 1 | 0.949005 |
| *Gr98a* | Branch-site Model A (ω = 1) | 2 | 16.922 | 0.732 | 0.2189 | 0.2189 | 0.268 | 1 | 1 | 0 | 0.2189 | 1 | 0 | 1 | 1 | -19599 | | | |

| | | | | | | | | | | | | | | | | | | | |
|---|---|---|---|---|---|---|---|---|---|---|---|---|---|---|---|---|---|---|---|
| *Gr98bcd* | Branch-site Model A | 2.08 | 23.611 | 0.728 | 0.16443 | 0.16443 | 0.267 | 1 | 1 | 0.004 | 0.16443 | 179.5489 | 0.001 | 1 | 180 | -12238 | 10.2 | 0.001404 | 0.046755 |
| *Gr98bcd* | Branch-site Model A (ω = 1) | 2.08 | 23.464 | 0.725 | 0.16312 | 0.16312 | 0.275 | 1 | 1 | 0 | 0.16312 | 1 | 0 | 1 | 1 | -12243 | | | |

### 2. SUPPLEMENTARY FIGURES

**Figure S1.** Distributions of spike responses to caffeine (10mM), lobeline (1mM), and sucrose (10mM) across all labellar sensilla (shown as a heatmap in **Fig. 2a**).

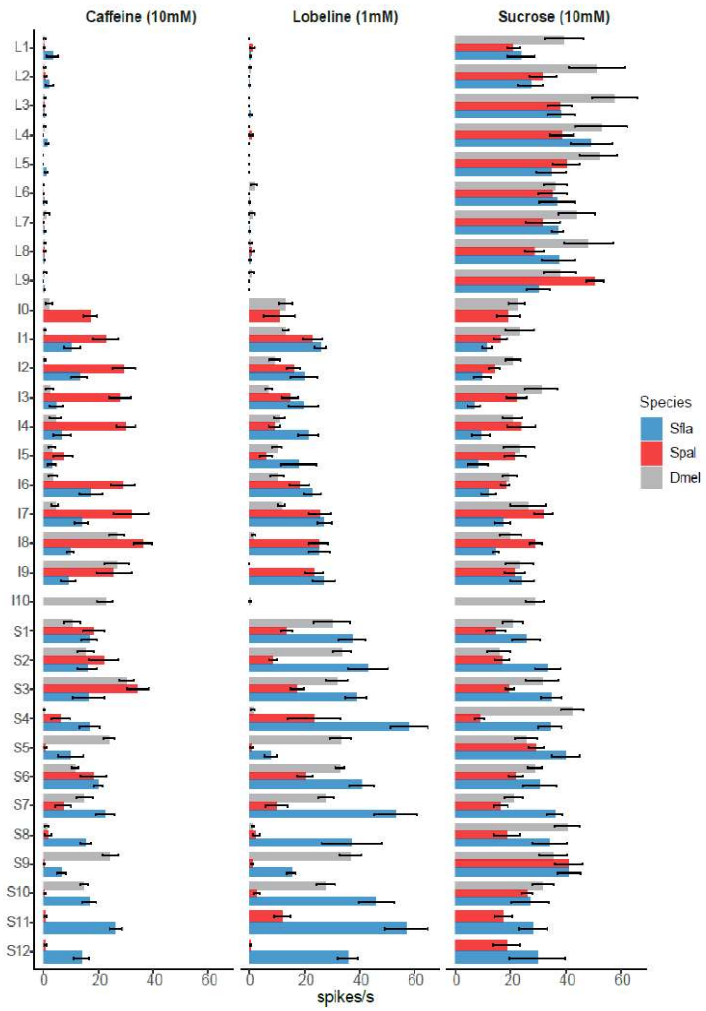

**Figure S2. Hierarchical cluster analysis to identify sensilla types on the labellum.** Based on Ward's classification method in the PAST program (Hammer et al., 2001) and using electrophysiological responses to caffeine, lobeline, and sucrose. Support values at the nodes were estimated with 100 bootstrap replicates.

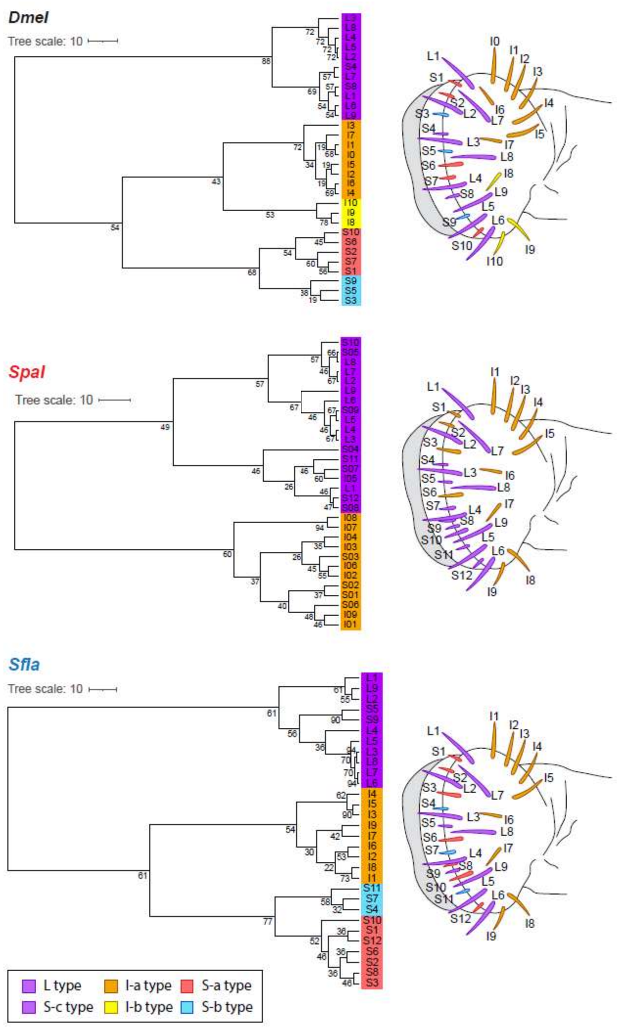

**Figure S3.** ***Scaptomyza*** **water cells do not respond to tricholine citrate the same as in** ***Drosophila melanogaster.*** Representative spike recordings from *S. flava* S5 sensilla (S-c type) at 3 concentrations of sucrose show an inverse relationship between the rate of sweet GRN firing (black bars) and water cell firing (gray bars).

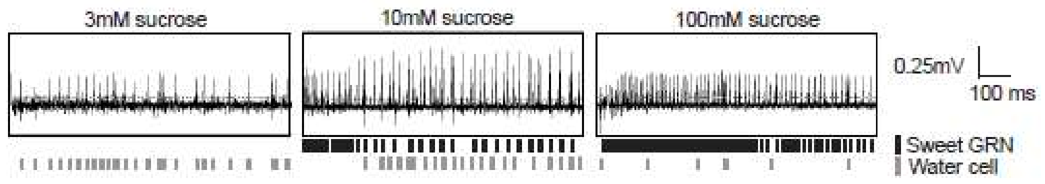

**Figure S4. Spike responses from all tested labellar sensilla.** (a) Illustration of sensilla representing each potential sensilla class that were recorded from in b, c, and d. Sensilla types marked with an asterisk are unlikely to be present in these species (based on responses) but were recorded from based on homologous length and location. Dose-dependent responses towards: (b) sucrose, lobeline, and caffeine; and (c) glucosinolates gluconasturtiin, neoglucobrassicin, and sinigrin. (d) Responses towards six glucosinolates (gluconasturtiin, glucotropaeolin, neoglucobrassicin, glucobrassicin, sinigrin, and glucoraphanin), all presented at a 10mM concentration. Error bars represent the standard error of the mean in b-c. (E) Spike responses from L type sensilla to a mixed solution of sucrose (5 mM) and sinigrin at two concentrations (1 and 10 mM), showing reduced sweet neuron firing in the presence of sinigrin in all three species.

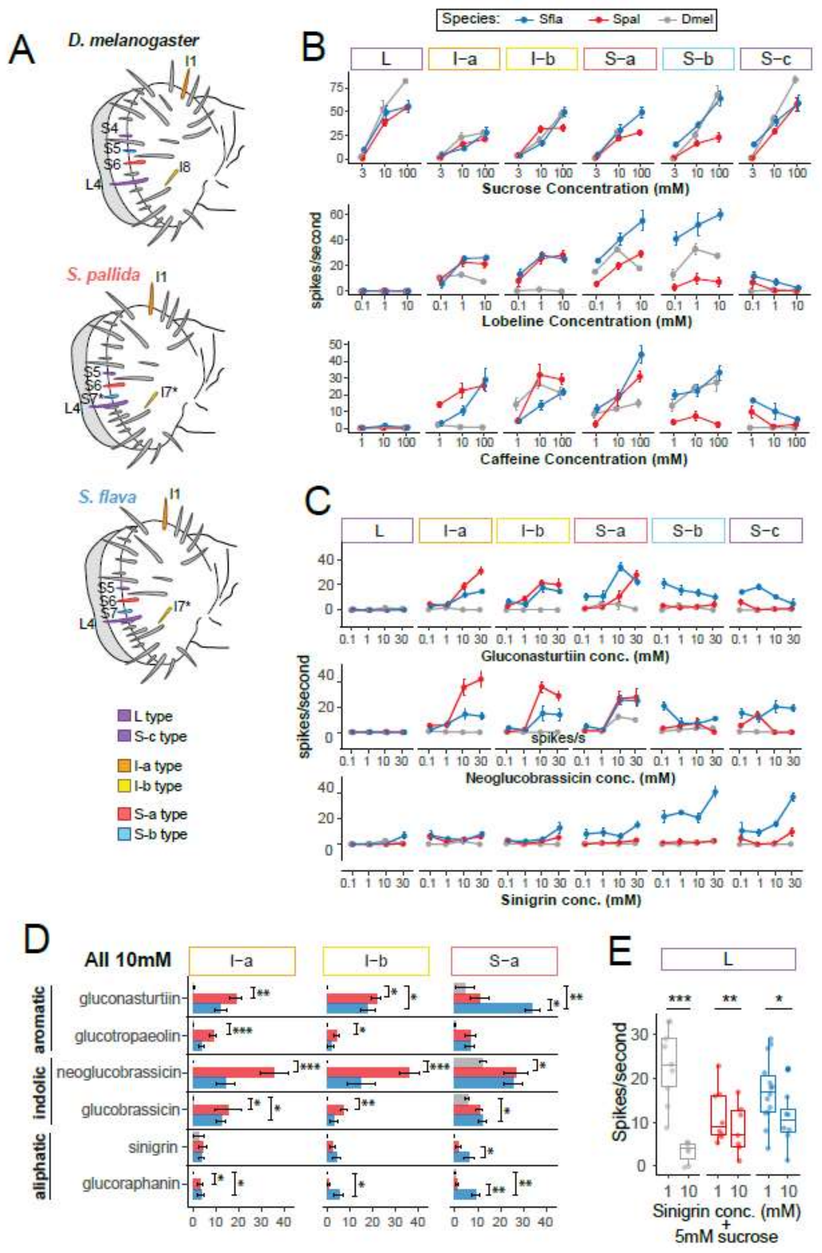
A
D. melanogaster
S. pallida
S. flava
L type
S-c type
I-a type
I-b type
S-a type
S-b type
B
Species: Sfla Spal Dmel
L
I–a
I–b
S–a
S–b
S–c
Sucrose Concentration (mM)
spikes/second
Lobeline Concentration (mM)
Caffeine Concentration (mM)
C
Gluconasturtiin conc. (mM)
Neoglucobrassicin conc. (mM)
Sinigrin conc. (mM)
D
All 10mM
aromatic
indolic
aliphatic
gluconasturtiin
glucotropaeolin
neoglucobrassicin
glucobrassicin
sinigrin
glucoraphanin
E
Spikes/second
Sinigrin conc. (mM)
+
5mM sucrose
